# Improving Cryo-EM Optimization Robustness with an Optimal Transport Loss Function for Noisy Images

**DOI:** 10.64898/2025.12.23.696001

**Authors:** Geoffrey Woollard, David Herreros, Minhuan Li, Pilar Cossio, Khanh Dao Duc

## Abstract

Many tasks in single-particle cryo-electron microscopy (cryo-EM), such as 2D/3D classification and homo/heterogeneous reconstruction, require optimizing model parameters to minimize the discrepancy between observed data and a forward model. The standard Mean Squared Error (MSE) loss function is computationally efficient but suffers from a non-convex rugged loss landscape, particularly for high-resolution heterogeneity inference. In this work, we investigate the practical utility of Sliced Wasserstein (SW) distances. We implement exact *W*_2_ estimators (inverse-CDF and greedy matching) of projections alongside a computationally efficient proxy based on the L2 norm of CDFs, a formulation akin to the sliced Cramér–von Mises distance. We establish the latter as a robust, fully differentiable workhorse for the cryo-EM forward model. We evaluate its performance against the MSE in joint inference tasks recovering pose, CTF parameters, and conformational heterogeneity. Our results demonstrate that SW significantly broadens the basin of attraction, enabling robust gradient-based optimization from distant initializations where MSE fails. Using a helical spiral toy model, we highlight how SW losses are sensitive to per-particle contrast, where background noise level miscalibration can induce geometric bias in the inferred structure. We show that this bias is manageable through a joint optimization strategy that treats background contrast as a learnable parameter. Finally, we validate the approach on a synthetic dataset using the Zernike3D framework, showing that the SW loss works and yields an accurate landscape representations, comparable with MSE. These findings establish SW as a powerful tool for navigating the rugged landscapes of cryo-EM forward model parameters.

**Synopsis:** The Sliced Wasserstein loss provides a smoother optimization landscapes than mean squared error for single particle cryo-EM joint inference of pose, CTF defocus and conformational heterogeneity. Estimating background contrast is essential to avoid biasing other parameters.

## 1 Introduction

Single-particle cryo-electron microscopy (cryo-EM) is a powerful technique for studying biomolecules at both high resolution and flexible conformations, in which 2D images of individual frozen molecules are observed with an electron microscope. A broad class of computational approaches have been developed to reconstruct the 3D structures of macromolecules from a large dataset of noisy images (Jensen, 2010; Toader *et al*., 2023; Donnat *et al*., 2022), spanning methods that recover a single static 3D density map to those that infer a continuous distribution of conformations. In a broad context, cryo-EM enables the investigation of biomolecular systems at near-atomic detail, allowing us to infer biophysical (structural, dynamical) properties. This inference is typically formulated as an optimization problem in which one seeks to recover a set of hidden (latent, *latens* in Latin) parameters - such as the specimen’s conformation, its ensemble distribution, the viewing direction (pose), microscope effects (e.g., the contrast transfer function, CTF), and the contribution of surrounding environment. These parameters are estimated by optimizing an appropriate objective (loss) function over the observed data, often together with prior information. The efficiency and robustness of this inference are therefore dictated by the choice of the loss function that measures the discrepancy between the the data and a forward model, where the loss is some distance or divergence in the “observed” space of 2D *n* × *n* pixilated images. Sometimes the loss is considered part of a statistical or probabilistic model/program, but here we consider the forward model distinct from the loss.

The mean squared error (MSE) loss remains the standard objective function in cryo-EM, with both advantages and limitations. It arises from the assumption that all pixels in the image are corrupted by independent and identically distributed Gaussian noise. A major benefit is computational efficiency: comparing two images with *n*^2^ pixels involves element-wise operations—subtraction, squaring, and summation—costing only 3*n*^2^ floating point operations. However, the resulting loss landscape for cryo-EM images is notoriously non-convex (Singer & Sigworth, 2020). By “loss landscape”, we mean the objective function in the space of latent variables (e.g., pose, heterogeneity, CTF, per-particle scale, …), comparing a fixed noisy observation to a noise-free simulation under a specific forward model simulator. While this landscape often contains a steep basin of attraction around the ground truth, it is plagued by local optima and flat plateaus beyond this narrow region, particularly at high resolution and for high levels of noise (Shi *et al*., 2025; Zelesko *et al*., 2020). This difficulty stems from a fundamental limitation: MSE treats images as statistical vectors, ignoring the underlying spatial topology. Because the calculation is invariant to pixel permutation, it fails to recognize when two features are spatially close but not overlapping at the given pixel discretization. This phenomenon is illustrated in Figure 1a: when a predicted signal (blue) is spatially separated from the target (gray), the MSE loss saturates at a constant value because there is no overlap between them. The resulting landscape exhibits vanishing gradients, providing no directional information to guide an optimizer toward the correct alignment. To some extent the point spread function (PSF, whose Fourier transform is the contrast transfer function, CTF) mitigates this issue by spreading the signal contained in each pixel outward, like ripples in a pond. However, as with ripples, the magnitude decays with distance from the source. As a result, gradient-based approaches using MSE (or other weighted Gaussian per-pixel likelihoods like coloured noise models) create a rugged loss landscape. To overcome some local minima, mature software implementations for standard cryo-EM data processing (Punjani *et al*., 2017; Scheres, 2012; Kimanius *et al*., 2021; Tang *et al*., 2007; Lyumkis *et al*., 2013; Grigorieff, 2016) rely on, for example, extensive grid sampling over particle orientations, or on search strategies like branch and bound or various heuristics in expectation maximization or other optimization strategies. Although some reconstruction methods employ gradient-based approaches (Punjani *et al*., 2017), these are typically applied only in the early stages of data processing and at low resolution. The difficulty of reconstructing volumes at high resolution by stochastic gradient descent has been examined in the setting of known poses by B. Toader and colleagues, who derived a preconditioner for faster convergence speed (Toader *et al*., 2025). If poses are known, the volume can also be parametrized by various implicit functions for rapid reconstruction with few gradient steps (Woollard *et al*., 2025). Other differentiable approaches often allocate a sizable budget of compute to initialization strategies of the orientation search to avoid getting trapped in local minima; for instance, cryoSPIN uses amortized inference with mixture models (Shekarforoush *et al*., 2024), while CryoDRGN (DRGN-AI) employs hierarchical pose search (Levy *et al*., 2025). Recent work using has demonstrated direct gradient-based optimization of atomic coordinates using individual images (Silva-Sánchez *et al*., 2025) and a physics based Hamiltonian, but did not yet achieve simultaneous inference of both pose and structure.

**Figure 1:**
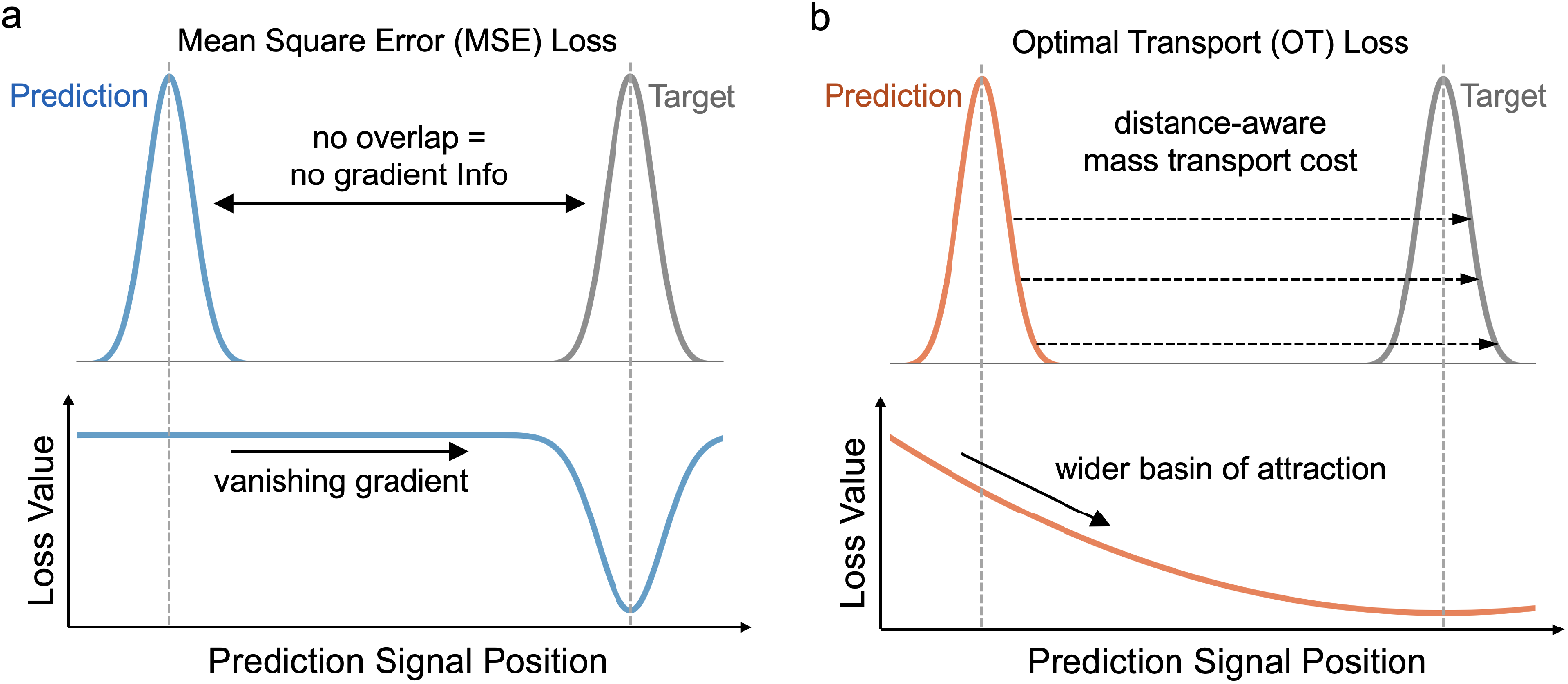
Conceptual comparison of MSE and OT loss. **(a)** MSE compares intensities locally. When the prediction model (blue) and target (gray) are separated, the error is saturated, leading to a flat loss profile with no gradient information. **(b)** OT loss explicitly measures the transport distance between signals (dashed arrows), creating a distance-aware, convex landscape that enables robust convergence even when the initial guess is far from the target.

An Optimal Transport (OT) loss between two images offers a compelling alternative by viewing images as a probability measure in image space *µ* ∈ 𝒫 (ℝ^2^), that is subsequently discretized into an image, and using displacement based distances that explicitly account for spatial locality/topology in the observed data space. In this framework, the distance between two images is framed as the physical “work” required to transport “mass” from one image to the other, rather than intensity overlap at fixed locations. As schematized in Figure 1b, this property can create a loss landscape with a wider basin of attraction that appears locally convex over a large region, providing a large basin of attraction. Unlike MSE, the OT loss provides informative gradients that scale with spatial displacement, enabling robust, end-to-end gradient-based inference even from distant initializations. OT losses have already been applied to cryo-EM in 2D classification (Rao *et al*., 2020) and 3D volume alignment settings with promising results due to wider basin of attraction (Riahi *et al*., 2023; Tajmir Riahi *et al*., 2025; Singer & Yang, 2024; Shi *et al*., 2025). OT was also used in the estimation of continuous conformational heterogeneity from simulated images using diffusion models (manifold learning): the distance kernel for affinities was build on top of a fast wavelet approximation of the *W*_1_ image to image distance, and deemed to be computationally tractable (Zelesko *et al*., 2020). However, replacing MSE with an OT loss in high-throughput end-to-end inference pipelines on large particle stacks that achieve competitive performance for single-particle cryo-EM poses further challenges. These include ensuring stable differentiability, improving computational feasibility, and scrutinizing statistical suitability (e.g., bias) under high noise.

In this study, we undertake such an effort, focusing on three problems: (i) inferring rigid body alignment between two 3D volumes, (ii) inferring structural and imaging parameters from individual noisy images, and (iii) inferring continuous conformational heterogeneity from a particle stack. For each of these tasks we use the Sliced Wasserstein (SW), a differentiable and computational efficient OT loss. While the Sinkhorn distance (Cuturi, 2013) offers an alternative computationally efficient route with better scaling for large images, it introduces an inherent entropic bias controlled by a regularization parameter that can be difficult to choose in practice and whose choice depends from case to case (e.g noise level). By contrast, SW provides a differentiable and efficient OT loss that avoids this entropic smoothing. We begin by demonstrating the robustness of the SW loss in a gradient-based 3D volume alignment task as discussed by previous work (Shi *et al*., 2025), showing it significantly outperforms MSE in recovering ground-truth poses from random initializations across diverse biomolecules. Transitioning to the problem of single-particle reconstruction from 2D projections, we systematically investigate differentiable SW implementations to balance computational cost with numerical accuracy. Specifically, we validate the use of the projected unweighted CDF L2 norm—related to the Cramér–von Mises distance (Cramr, 1928; Anderson, 1962)—as a highly efficient proxy for the 2D Wasserstein distance. This formulation avoids the computational overhead of quantile inversion while preserving the smooth optimization landscape. We validate this efficient implementation in a gradient-based joint-inference setup that simultaneously estimates local per-image latent variables, including conformational heterogeneity, rotation, translation, and CTF defocus. Next, we identify image noise as a key factor in OT loss performance using a helical spiral toy example, constructed to be pathological to MSE. We show that accurate calibration of background noise is critical for the SW objective to avoid biased estimates; and crucially, we demonstrate that this calibration can be achieved by including the noise level as a differentiable parameter to be jointly optimized. Applying these insights, we demonstrate the feasibility and advantage of SW in fully differentiable gradient-based pipelines. We show that SW loss successfully infers heterogeneity over an entire dataset in an established deep learning framework: Zernike3D (Herreros *et al*., 2023). Finally, we provide suggestions for applying the OT loss to real data where noise is a hidden variable, and invite methods developers to engage with the existing body of theoretical results on OT losses. To this end, we release a modular software implementation supporting JAX and PyTorch backends to enable the community to easily adapt the SW loss into existing pipelines.

## 2 Methods

### 2.1 Forward model and experimental setup

We utilized two differentiable forward models to map atomic coordinates to single-particle cryo-EM images: the JAX-based `cryoJAX` package (O’Brien *et al*., 2025) (version 0.5.0rc4.dev166+g383e39ab5) and a custom differentiable projector implemented in PyTorch. By validating our loss functions against forward models in both JAX and PyTorch, we explicitly demonstrate that our modular implementation is robust and interoperable across the two dominant deep learning backends. Further implementation details are provided in Extended Methods A.1.

### 2.2 Noise model and parameterization

Simulating realistic noise is critical for evaluating inference robustness, particularly for transport-based losses which are sensitive to mass conservation. While many computational studies employ additive Gaussian white noise, real cryo-EM data is physically generated by electron counting events, which approach a Gaussian distribution in a high count regime. Counts are also always non-negative. We therefore model the observed noisy image *y* with Poisson shot noise of a contrast scaled CTF abberated projection, *ŷ*, using cryoJAX’s intensity mode which gives rise to an image that is everywhere positive. The shot noise is parameterized by a total dose *d* (electrons per pix^2^) and a background contrast *b* that is relative to the scale of *ŷ*:

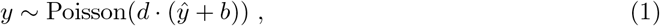

We chose this Poisson formulation over the more popular Gaussian model to better capture the signal-dependent variance inherent in low-dose microscopy, the non-additive nature of shot noise whose SNR depends on both *d* and *b*, and the positive co-domain of electron counts in the observation. To assist readers familiar with Gaussian signal-to-noise ratios (SNR), we provide a theoretical comparison based on KL divergence in Extended Methods A.2 and a visual correspondence in Appendix Figure A.1. Crucially, the performance of the Optimal Transport loss is heavily dependent on the accurate calibration of this background noise level. We explore the implications of this sensitivity and demonstrate a strategy for joint noise estimation in Section 3.3. In practice, once the dose *d* and background level *b* are known (or estimated), we explicitly calibrate the noise-free forward model output as *y*′ = *d*(*ŷ* + *b*) before computing the loss against the noisy observation *y*. If the observation has been normalized, we backtransform it to match the same scale as *y*′, as explained in Section A.1.5, which assumes the noise model in Eq. (1).

### 2.3 Optimal Transport Loss

Exact calculation of Optimal Transport distance is known to scale poorly with problem size, becoming computationally prohibitive even for standard 2D images (many seconds) or 3D volumes. To achieve the goal of an optimal transport loss that is computationally feasible and differentiable, we adopted the Sliced Wasserstein (SW) framework. The SW distance simplifies the problem by projecting high-dimensional distributions onto random 1D lines (Bonneel *et al*., 2015), where the transport cost can be computed efficiently. We explored three differentiable implementations for this 1D calculation: (1) the norm between inverse cumulative distribution functions (iCDF), (2) the norm of the cumulative distribution functions (CDF), and (3) a “greedy match” (GM) approach that performs a single pass through the marginals without backtracking. As a ground-truth baseline for validation, we utilized the exact *W*_2_ calculation provided by Python OT (POT) library `ot.emd2` (Flamary *et al*., 2021; Flamary *et al*., 2024). Numerical tests quantifying the accuracy of our approximations against this baseline are detailed in Appendix section B. Throughout this work, we assume all images are non-negative to satisfy the requirements for probability measures. As explained in the Section 2.2, we achieved this for simulated images with cryoJAX’s intensity mode.

#### 2.3.1 2D Earth Mover’s Distance (Baseline)

For this exact baseline, we defined the cost matrix as the squared Euclidean distance between pixels, normalized to a maximum of 1.0 for numerical stability and use POT’s `ot.emd`, which uses a network simplex solver with a large computational cost: *O*(*n*^6^) for an *n* × *n* image. We utilized the analytic gradient hooks provided by the POT backend, which compute gradients using the optimal transport plan and did not unroll the optimization loop for differentiation. See Appendix A.3 for implementation details).

#### 2.3.2 Sliced Wasserstein

To overcome the computational bottleneck of the exact Earth Mover’s Distance, we adopted the SW distance (Bonneel *et al*., 2015). The SW distance approximates the Wasserstein by projecting the high-dimensional 2D marginals onto 1D lines (slices), computing the 1D Wasserstein distance along each slice, and averaging the results. We investigated two distinct strategies for performing these projections and computing the subsequent transport cost, which are summarized below. Ssee Appendix A.4 for more implementation details.

##### Method 1: Sparse Projections via Coordinate Rotation

We treated the image/volume as a collection of weighted masses at specific coordinates. Instead of interpolating the grid, we rotated the coordinate vectors of the pixels and projected them onto the 1D line by indexing the transformed coordinates. This approach enables sparse subsampling of pixels, which prevents averaging out high-frequency noise but significantly reduces computational load. Because this method yields discrete, non-uniform supports on the 1D line, we differentiably sorted each space and then computed the Wasserstein distance using a differentiable re-implementation of POT’s *Greedy Matching* algorithm (*SW*_GM_). This algorithm performs a single-pass greedy assignment of mass.

##### Method 2: Dense Projections via Radon Transform

In this approach, we project the full image (not implemented for volumes) density onto a set of fixed angles on the circle —effectively computing the Radon transform. We implemented this differentiably by rotating the grid using linear interpolation and summing pixel intensities along the vertical axis. Once projected to 1D distributions, we computed the transport distance using two formulations:

- **Greedy Matching (***SW*_**GM**_): After the Randon transform the space is already sorted along the equidistance strip of projected pixels. We computed the the exact 1D Wasserstein distance *W*_2_ via greedy matching (as above). To clarify, there was no random pixel subsampling, and the space is already sorted.
- **Inverse CDF (***SW*_**iCDF**_): Following the analytical solution for 1D *W*_2_ (Remark 2.30 in (Peyré & Cuturi, 2019)), we integrated the squared difference between the inverse cumulative distribution functions of the projected marginals.
- **CDF Norm (***SW*_**CDF**_): As a computationally efficient proxy, we computed the direct L2 norm between the cumulative distribution functions (CDF) of 1D projections. This formulation is closely related to the unweighted Cramér–von Mises distance (Cramr, 1928; Anderson, 1962) rather than the Wasserstein distance. We found this simpler metric yielded sufficient numerical agreement for optimization purposes while avoiding the interpolation costs of the inverse CDF method. For the sake of notational consistency, we also refer to this metric as SW throughout the manuscript, as it serves as a functional proxy that shares the similar topological properties and in our experiments the same global minimum as the canonical Wasserstein distance.

### 2.4 Zernike3D

Zernike3D (Herreros *et al*., 2023) is an approach previously introduced in the field to estimate diverse structural states from a set of particle images in a cryo-EM dataset. In brief, starting from a 3D reference state (e.g., a consensus volume), the method estimates the vector field (parametrized by the Zernike3D basis) wrapping the reference geometry towards a target state captured in a 2D image. The coefficients for the per-image vector field are estimated by a deep neural network conditioned on the image. The global parameters of the deep neural network are optimized through an image-to-image loss that compares the noisy image to the clean CTF aberrated and vector-field-warped simulated image. More details are in extended methods section A.6.

## 3 Results

### 3.1 Robust gradient-based 3D volume alignment

Before addressing the challenging task of multi-parameter inference from noisy 2D images, we begin with a simpler problem that requires estimating only a single parameter in a noise-free landscape: the 3D rigid-body rotational alignment between a density volume to itself. For these numerical experiments, we utilized the sparse Greedy Matching implementation (*SW*_GM_). Since 3D volumes are sparser (more empty space) compared to dense 2D projections, the coordinate-based greedy matching approach is computationally efficient in this setting. We compiled a dataset of 13 diverse biomolecules (see Appendix A.5) and, for each system, generated 100 random initial poses sampled uniformly within 90-degree geodesic distance from the ground truth. We then performed gradient-based optimization to align each volumes to itself using either SW or MSE loss. Our approach for this task is related to Algorithm 1 in (Riahi *et al*., 2023) but does not require or make use of an optimal transport plan for correspondences. We summarized the convergence performance by analyzing the cumulative distribution function (CDF) of the rotational errors before and after optimization (Figure 2a). An ideal optimization would drive all random initializations to the ground truth (0 degrees error), resulting in a CDF that resembles a step function at the origin. We quantified the convergence performance using the Area Under the Curve (AUC) of the CDF, where an AUC of 1.0 represents perfect convergence from all initializations. As summarized in Figure 2b, the SW loss demonstrates superior robustness. Across all 13 test systems, the final AUC values for SW are tightly clustered near 1.0, indicating that the loss landscape is sufficiently smooth to guide the optimizer to the global minimum even from large initial misalignments. Conversely, the MSE loss performance is highly variable; the wide spread of AUC values reflects the non-convex nature of the voxel-wise comparisons, where the optimizer frequently stalls in local minima. We note, however, that this distinct advantage was less apparent when aligning 3D coordinates to 2D projection images (data not shown), suggesting that the distribution of mass in 3D space plays a key role in the robustness of the transport distance. This motivates the need for a rigorous evaluation of SW implementations in the context of 2D projections.

**Figure 2:**
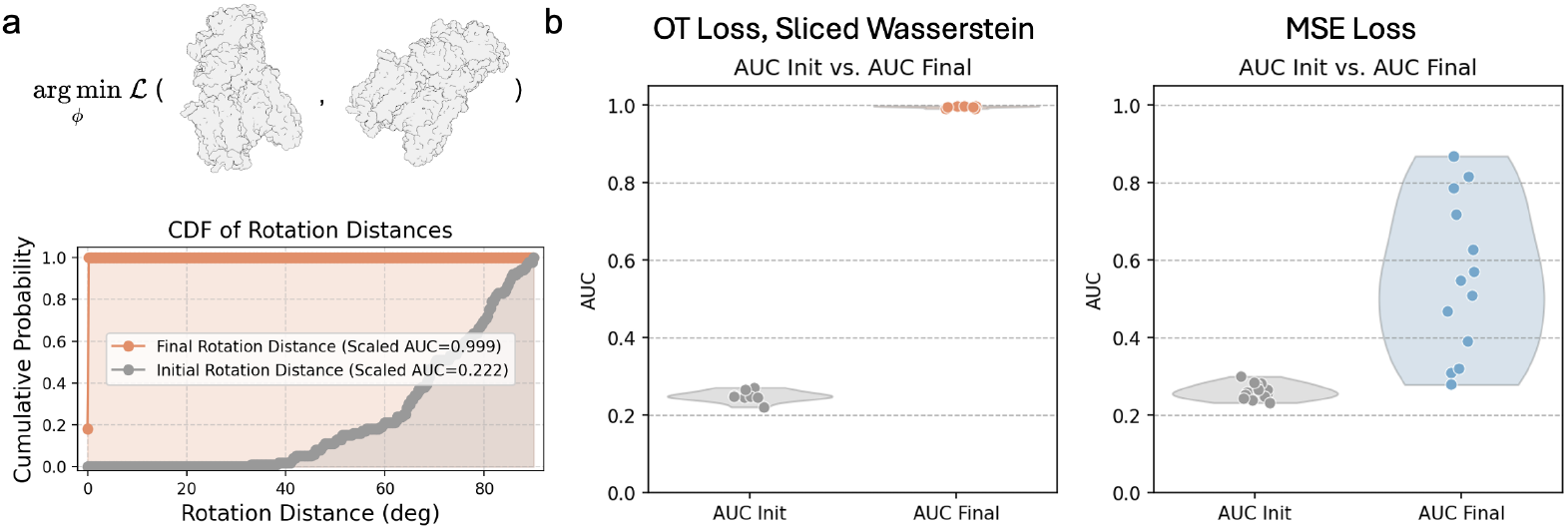
Optimal Transport (SW) loss enables robust gradient-based 3D volume alignment compared to MSE. **(a)** Top: Schematic of the 3D alignment task, where a source volume is rotated (*ϕ*) to match a target volume by minimizing the loss L. Bottom: Representative performance on a single system. The Cumulative Distribution Function (CDF) of rotation errors is plotted for 100 random initializations (Initial, gray) and their optimized final states (Final, orange). A steep rise near zero indicates successful convergence. We quantify this performance using the normalized Area Under the Curve (AUC), where AUC ≈ 1.0 indicates perfect convergence from all initializations. **(b)** Summary of alignment performance across 13 diverse biomolecular systems (see Appendix A.5). Each point represents the AUC for a specific system before and after optimization. The SW loss (left) yields a tight cluster of AUCs near 1.0, indicating consistent recovery of the ground truth pose across all systems. In contrast, the MSE loss (right) shows high variance and lower AUCs, reflecting frequent entrapment in local minima.

### 3.2 Efficient Sliced Wasserstein enables robust joint optimization on a single-particle image with known background contrast

While the 3D alignment result confirms the favorable basin of attraction of SW in 3D space, single-particle cryo-EM requires inferring structural and imaging parameters from 2D projections. This projection step introduces significant computational overhead and spatial correspondence ambiguity, not to mention microscope effects and noise. Therefore, to enable the use of OT in high-throughput cryo-EM pipelines, we next sought to identify a SW implementation that balances numerical accuracy with computational efficiency.

We constructed a representative joint inference using a forward model implemented in cryoJAX. The goal is to infer the simulator’s input parameters, along with a 1D conformational collective variable (CV) that acts directly on coordinates, from a single noisy image. We used the PDB structure 1UAO of Chignolin, a 10 amino acid artificial peptide/mini-protein, and split it into two sets of atoms by a plane crossing the center of mass. We defined the split distance between these sets as the CV (*x*), alongside the standard rigid-body pose parameters: 3D rotation (*R*, inference done in quaternion representation), 2D translation (*t*), and CTF defocus (*f*) (Figure 3a). Our goal is to infer these from the single-particle image. We generated a ground truth image with the protein domains centered (*x* = 0, *t*_*x,y*_ = 0), unrotated (*R*_*ϕ,θ,ψ*_ = 0), and at a defocus of *f* = 2000 Å—the resulting noise-free image ranged ~ 0.5 − 1.3. Then Poisson shot noise was added (*d* = 100, *b* = 1) to produce a noisy observation. To simulate a difficult inference scenario far from the basin of attraction, we initialized the prediction with sensible perturbations: the domains were separated by 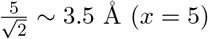 (*x* = 5), the particle was rotated and translated ( *R*_*ϕ,θ,ψ*_ = 5°, *t*_*x*_ = *t*_*y*_ = 5 Å), and the defocus was offset to *f* = 3000 Å.

**Figure 3:**
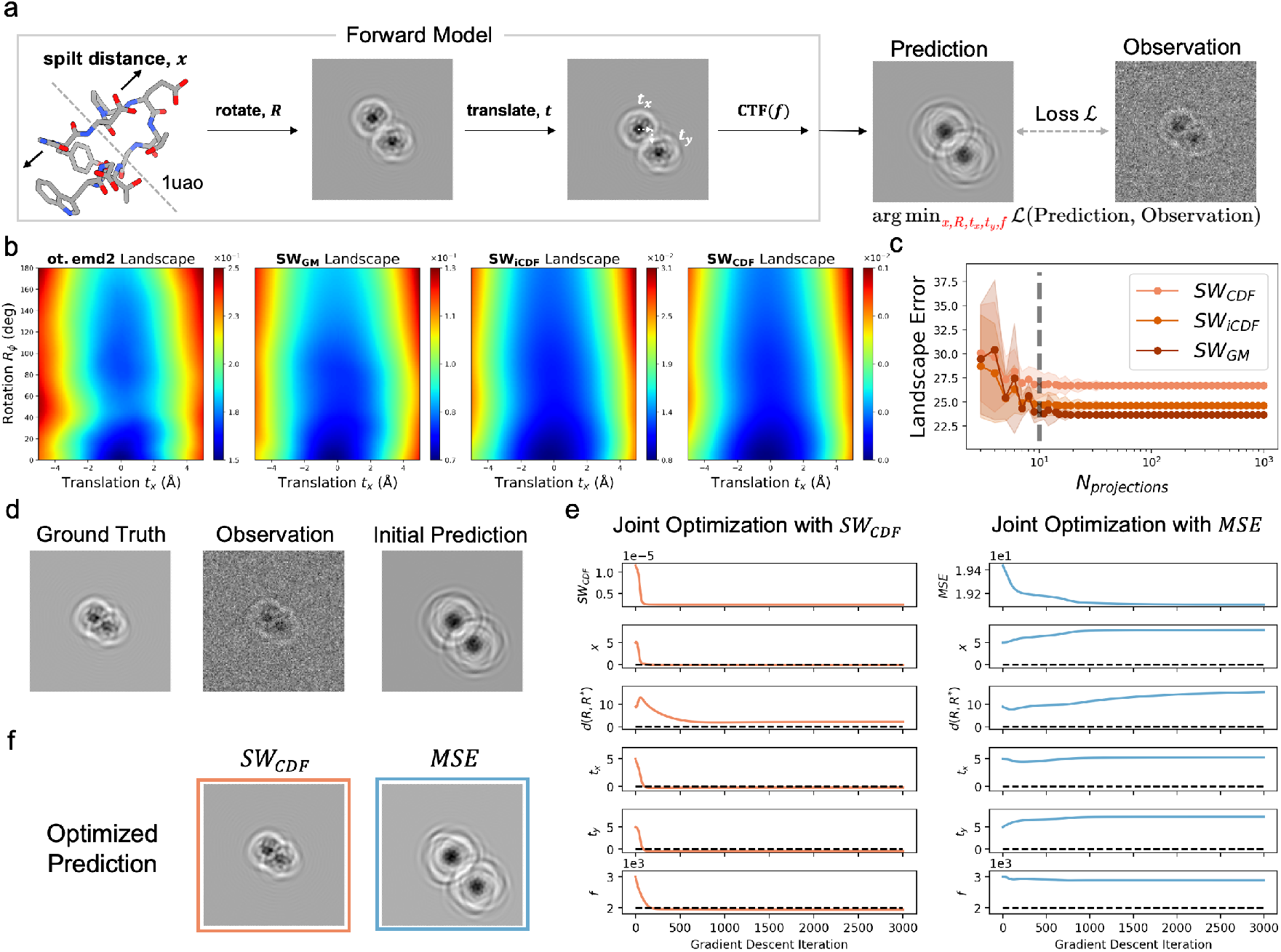
Establishing computational feasibility and performance of Sliced Wasserstein (SW) loss in a joint cryo-EM inference task. **(a)** Schematic of the forward model used for joint inference. A protein structure (PDB: 1UAO) is split into two domains, with their separation distance defined as the conformational variable (*x*). The model generates a 2D projection based on this conformation, 3D rotation (*R*), 2D translation (*t* = (*t*_*x*_, *t*_*y*_)), and CTF defocus (*f*), corrupted by Poisson noise parameterzied by dose and background. **(b)** Loss landscapes over rotation angle *R*_*ϕ*_ (about axis (1, 1, 1)) and translation *t*_*x*_(ground truth at 0,0) comparing the exact Earth Mover’s Distance baseline (`ot.emd2`) with three differentiable SW implementations (CDF, iCDF, GM) using *N*_projections_ = 1000. All approximations correctly recover the smooth basin of attraction centered at the optimum. **(c)** Analysis of approximation error relative to the `ot.emd2` baseline as a function of the number of projections (*N*_projections_). The error plateaus near *N*_projections_ = 10, identifying a computationally efficient operating point. A ± 1 std envelope is shown, where the variance was computed over distinct sets of different, but still evenly spaced, projection angles. **(d)** Setup for the gradient-based joint inference experiment comparing the efficient *SW*_CDF_ implementation (*N*_projections_ = 10) against the standard MSE loss. Initial predictions were perturbed from the ground truth across all latent variables (*x, R, t, f*), and optimization was performed separately for each loss function. **(e)** Optimization trajectories comparing *SW*_CDF_ loss (orange) and MSE loss (blue). *SW*_CDF_ successfully drives all parameters to the ground truth, whereas MSE fails to converge due to local minima. **(f)** Final reconstructed images. *SW*_CDF_ recovers the true projection features, while the MSE result remains stuck near the initialization.

We evaluated three fully differentiable SW implementations—CDF-based, inverse-CDF (iCDF), and greedy matching (GM)—by comparing their loss landscapes, in translation and rotation, against a baseline calculated using the computationally expensive exact *W*_2_, computed with POT’s Earth Mover’s Distance (`ot.emd2`) with squared Euclidean cost. As shown in Figure 3b, when the background level is correctly calibrated, all three approximations yield smooth, seemingly convex landscapes on the rotation angle (*R*_*ϕ*_) and translation (*t*_*x*_) plane that closely resemble the exact baseline, with the global minimum correctly located at the ground truth (*R*_*ϕ*_ = 0°, *t*_*x*_ = 0 Å). A critical parameter for SW is the number of random projections (*N*_projections_), which scales linearly with computational cost. We quantified the approximation error of the SW landscape relative to the exact OT baseline as a function of *N*_projections_. We observed that the error decreases rapidly and plateaus at approximately *N*_projections_ = 10 for all three implementations (Figure 3c). Consequently, for subsequent experiments, we selected the CDF-based implementation with *N*_projections_ = 10, unless otherwise stated. While formally constituting an unweighted variant of the Cramér–von Mises distance proxy rather than the exact Wasserstein distance, it offers a sufficiently accurate approximation while being the simplest and fastest to compute.

With the efficient *SW*_CDF_ (*N*_projections_ = 10), we challenged the loss function in a fully differentiable joint inference task. We sought to recover all hidden parameters-conformation (*x*), pose (*R, t*), and CTF defocus (*f*)—simultaneously using gradient descent with per-parameter step sizes to account for differently scaled units (Table 2). Note that in this experiment, we assume the noise level is known and calibrated. A more detailed discussion on the noise will be covered in section 3.3. We ran the joint optimization to minimize the discrepancy between the predicted projection and the noisy observation, comparing the performance of the standard MSE loss against our *SW*_CDF_ loss. The optimization trajectories in Figure 3e reveal a stark contrast in performance dynamics. The MSE loss failed to converge due to the roughness of the pixel-wise landscape; indeed, the optimization not only got trapped but caused several parameters to diverge further from the initialization. In contrast, the *SW*_CDF_ loss trajectories show a rapid and stable descent toward the ground truth (Figure 3e, orange lines). We show the accuracy of the optimization trajectory end points in Table 1. While the MSE rotational error worsened from the initial 8.9° to 15.4°, the *SW*_CDF_ loss achieved a final rotational error of just 2.3°. Similarly, SW reduced the defocus residual from 1000 Å to 50 Å. Notably, Table 1 demonstrates that the highest precision is achieved by leveraging the complementary strengths of both functions. By employing a sequential strategy —using SW to navigate the landscape to point closer to optimal, then switching to MSE for local refinement—we surpassed the performance of SW alone. As shown in the “SW+MSE” row of Table 1, this hybrid approach further reduced the rotational error to 1.0° and the defocus residual to a negligible 10 Å.

**Table 1:**
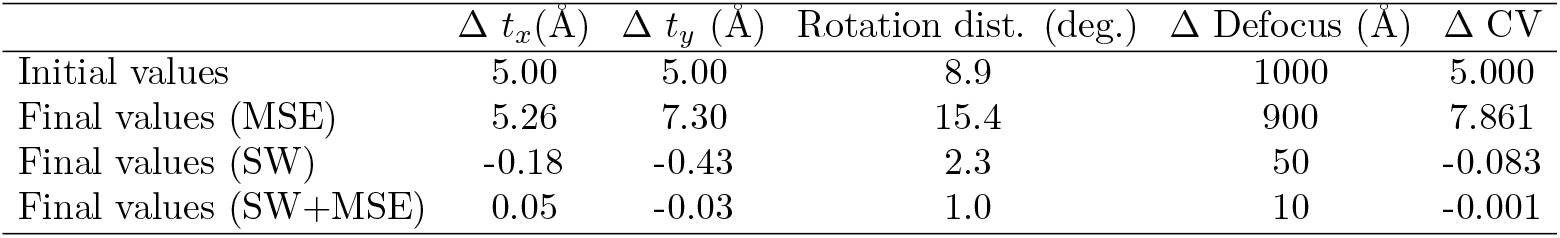
Joint inference parameter. Residuals (Δ) are defined as the respective parameter value minus the ground truth in the noisy reference image: 5 Å x and y translation, 0.3 *µ*m defocus, and a CV value of 0. For rotation, we show the geodesic distance with respect to the ground truth identity rotation—a perfect rotation distance would be 0 degrees.

**Table 2:** Preconditioner scaling values used for different loss functions.

| Loss | CV | Defocus | Rotation (quaternion) | Translation |
| --- | --- | --- | --- | --- |
| MSE | $1 \cdot 10^{-1}$ | $1 \cdot 10^4$ | $1 \cdot 10^{-4}$ | $1 \cdot 10^{-1}$ |
| $SW_{\text{CDF}}$ | $5 \cdot 10^5$ | $1 \cdot 10^{10}$ | $3 \cdot 10^2$ | $1 \cdot 10^5$ |

We further analyzed the computational cost of this substitution. In our test loop, a single iteration, including forward pass and gradient update, with the MSE loss took 0.0027 s, while the *SW*_CDF_ loss took 0.0122 s (CPU on a Linux-based computer cluster). For context, the forward model evaluation alone without loss or gradient computation takes 0.0011 s, 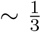 the time of the gradient descent iteration with MSE. While the SW calculation is approximately 4.5× slower than MSE, this overhead must be viewed in the context of the total computational budget. As forward models evolve to include complex transformations, such as neural radiance fields (NeRF) (Zhong *et al*., 2019; Woollard *et al*., 2025; Qu *et al*., 2025), Gaussian splatting (Chen & Ludtke, 2021; Chen & Ling, 2025), Langevin dynamics (Silva-Sánchez *et al*., 2025), or AlphaFold-like generative backbones (Raghu *et al*., 2025), we speculate that the cost of generating the prediction (on the fly each iteration) may rise by orders of magnitude —at least 10× the time of the rest of the forward model seems reasonable. In such regimes, the relative cost of the loss function is not the computational bottle neck, making the robust convergence properties of SW a highly favorable trade-off for heterogeneity inference, particularly in any task where the MSE basin of attraction is narrow relative to SW.

### 3.3 Background noise calibration is essential to avoid bias

In the previous joint inference experiments, we assumed the noise level was perfectly calibrated. However, in empirical cryo-EM settings, the background noise level (contrast) is often an unknown latent variable. To investigate the interplay between noise calibration and structural inference, we designed a “helical spiral” toy model that undergoes a complex expansion/contraction motion along a conformational collective variable —a different CV than in the joint inference experiment; details in Extended Methods A.1.4. This system is designed to be a rough landscape where the ridges of the spiral move in and out of registration, creating severe multiple minima for pixel-wise comparisons like MSE.

We analyzed the loss landscapes for both *SW*_CDF_ and MSE by comparing a fixed noisy observation (Ground Truth: *CV* = 4, *b* = 100) against predictions with varying conformal states and background levels. As shown in Figure 4 (middle right), the MSE landscape is riddled with local minima, possessing only a narrow “funnel” of attraction near the ground truth. While the MSE global minimum is statistically robust—locating *CV* = 4 with low variance even with high shot noise—finding this minimum via gradient descent fails from a distant initialization. In contrast, the *SW*_CDF_ landscape (Figure 4, middle left) is a bowl, avoiding the local minima that plague the MSE. However, this smoothness comes with a specific sensitivity. Because OT is fundamentally a mass-moving cost, it is sensitive to the total intensity of the image. If the background level is misestimated (e.g., underestimated), the SW loss drives the spiral to physically distort (stretch or squeeze) to compensate for the missing or excess mass. This creates a geometric bias where errors in contrast estimation propagate directly into errors in structural inference. While expected, if not controlled this could lead to overly stretched or squeezed shape that is very different than artefacts (noise dust, spikes, blobs) under an MSE loss.

**Figure 4:**
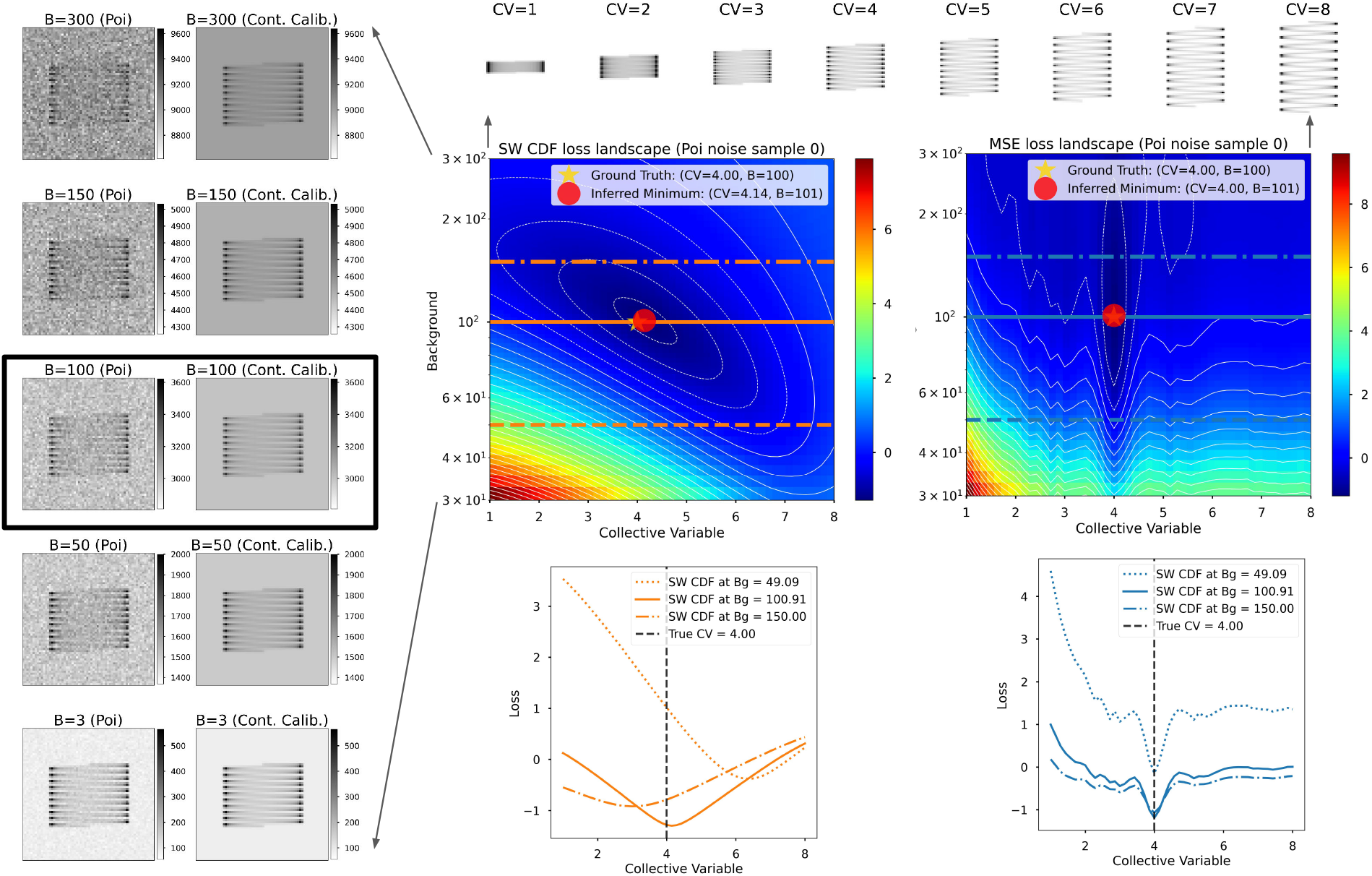
Accurate background noise contrast calibration is essential to prevent statistical bias in Sliced Wasserstein inference. **(Top)** A helical spiral toy model parametrized by a 1D stretching/squeezing conformational collective variable (CV). **(Middle)** Loss landscapes comparing a fixed noisy observation (Ground Truth: *CV* = 4, *b* = 100) against simulated projections with varying CV and background (*b*) levels. The landscapes are normalized by Z-score for visual comparison. The SW landscape (left) is smooth and jointly convex. However, the optimal CV depends on *b*; underestimating the background level causes the model to “stretch” the spiral (higher CV) to conserve mass. In contrast, the MSE landscape (right) is plagued by local minima but is statistically unbiased; the optimal CV remains at 4.0 regardless of the background level. **(Bottom)** 1D slices through the landscapes at fixed background levels (*b* ≈ 50, 100, 150, indicated by dashed lines). The MSE slices show a narrow and deep local minima that guide gradient descent to the true CV value if initialized within ±0.5. The SW slices are a smooth bowl but shifted, illustrating that accurate background inference is a prerequisite for unbiased structural recovery. Note that the MSE and SW losses have been square rooted on the middle and bottom panels to illustrate trends.

Crucially, however, the landscape reveals that this bias is resolvable. Because the *SW*_CDF_ landscape is seemingly jointly convex with respect to both the conformational variable and the background level (within a few fold to the true background level), we can treat the background as a learnable parameter. As illustrated in Figure 4, the joint global minimum exists effectively at the ground truth (CV≈4.14, b≈101). This implies that while SW requires accurate noise calibration to avoid bias, it uniquely provides the gradient information necessary to perform that calibration on-the-fly. We further quantified this behavior across 100 random noise instantiations at various noise levels (Appendix C), confirming that while higher noise levels increase the variance of the bias, joint optimization would be feasible.

### 3.4 Dataset distribution learning with SW loss in Zernike3D pipeline

While the previous experiments demonstrated the theoretical robustness of *SW*_CDF_ on single-image optimization and toy models, practical cryo-EM reconstruction requires aggregating signal from thousands of noisy images to resolve continuous heterogeneity. Here, we integrate the *SW*_CDF_ loss into Zernike3D, a previously published end-to-end gradient based method.

The dataset comprises an open-to-close trajectory of the adenylate kinase protein (PDB entry 4AKE) simulated using Normal Mode Analysis with HEMNMA (Harastani *et al*., 2021). To simplify the simulated landscape, only two modes were excited and uniformly sampled, resulting in a ground-truth landscape with a straight-line shape. The sampled structures were then further processed with Xmipp (de la Rosa-Trevin *et al*., 2013) inside Scipion (de la Rosa-Trevin *et al*., 2016) to generate a set of 500 CTF corrupted projections with the rotation distribution uniform on SO(3).

#### 3.4.1 Performance on noise-free images

Our first experiment compared the *SW*_CDF_ loss function proposed in this work with MSE for motion estimation directly from images. Therefore, the 500 simulated images were subjected to two independent Zernike3D analyses, each using the previous two loss functions to optimize the Zernike3D coefficients. The results of this experiment are summarized in Figure 5. In the figure, we present the two Zernike3D landscapes derived from the optimized Zernike3D coefficients, the principal curve that crosses the landscape, and some representative conformational states recovered from the motion fields. Although the two Zernike3D landscapes have adequately captured the overall shape of the ground truth space, it is evident that the *SW*_CDF_-optimized space consistently concentrates points along the landscape’s principal curve. Therefore, thanks to the *SW*_CDF_ loss, it is possible to obtain a more accurate representation of the expected landscape with an intrinsic dimensionality closer to one. We also used the rank based method of information imbalance previously used in (Jeon *et al*., 2024) to quantitatively compare the inferred deformation fields (encoded as Zernike3D coefficients) to the ground truth heterogeneity (encoded as normal mode coefficients), and found that using the *SW*_CDF_ the estimated heterogeneity is slightly more accurate (Appendix Figure A.5 and Appendix D).

**Figure 5:**
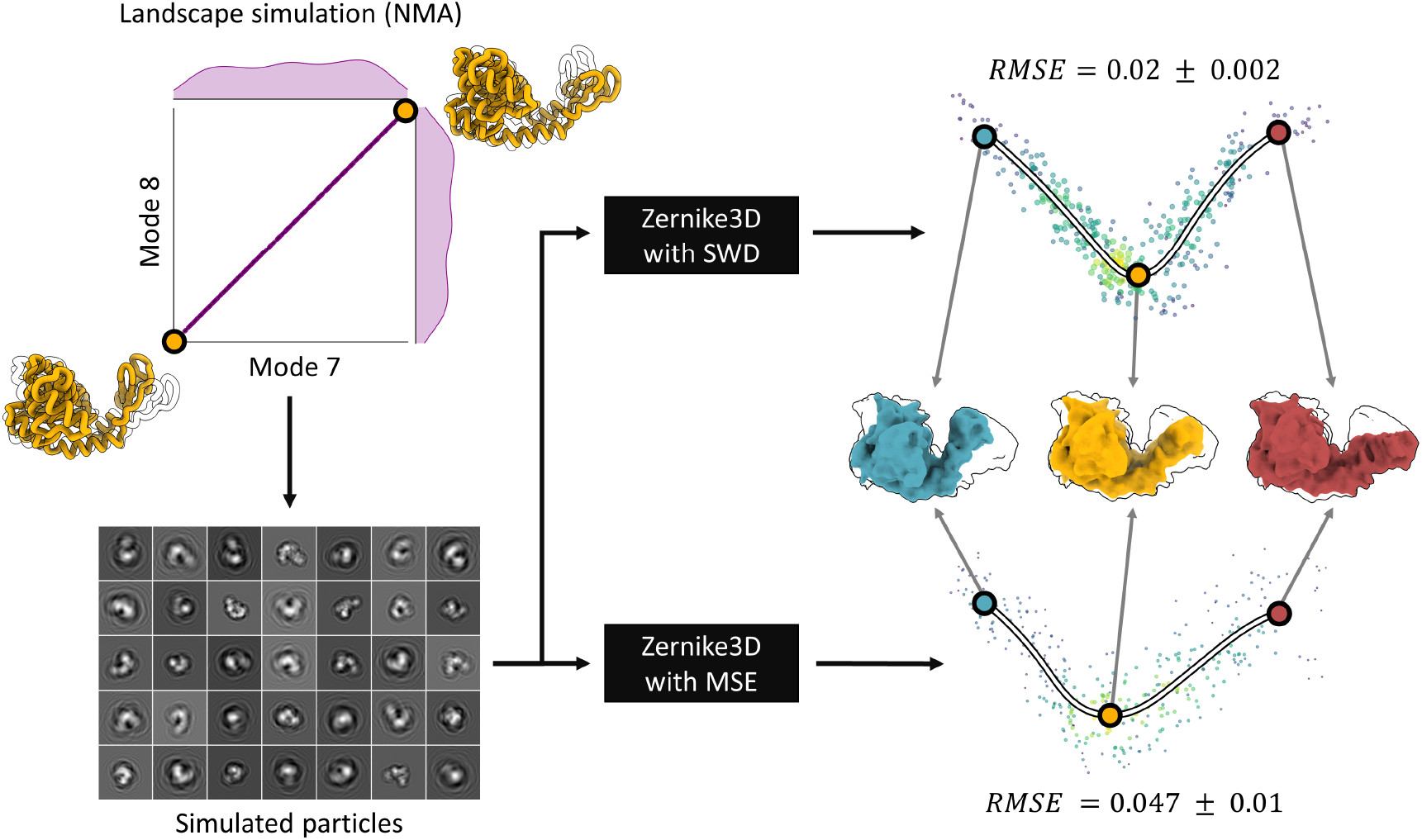
Zernike3D comparison of MSE and SWD. Schema of the adenylase kinase dataset simulation process, followed by the Zernike3D results obtained from both the MSE and SWD losses. Although both loss functions yielded a good approximation of the ground-truth landscape, the SWD distance tends to reduce the spread of points, leading to a representation closer to the simulated space with intrinsic dimension one.

#### 3.4.2 Effect of noise and contrast calibration in experimental data

From the previous experiment, a question remains on how the Sliced Wasserstein Distance would perform when comparing simulated projections of a volume against experimental images. When using the SW, it is essential to distinguish between two factors: (1) the noise and (2) the contrast mismatches resulting from variable ice thickness and image normalization during a processing workflow (see also helical spiral results in section 3.3).

To better assess each effect independently, we first focus on the noise effect. We conducted an ablation test in which we progressively corrupted the CTF-corrected images analyzed in the previous section with noise, up to an SNR comparable to that expected in experimental cryo-EM images. The noise added to the images followed a Poisson distribution, followed by a normalization step to ensure the images’ backgrounds had zero mean and unit standard deviation. It is important to note that, thanks to this controlled scenario, we are ensuring consistent background normalization across all images, which would rarely be the case in an experimental dataset.

The noisy images were then subjected to the Zernike3D analysis with the *SW*_CDF_ loss, with positive and negative mass separately normalized and matched to each other in two additive terms. We termed this loss SWD. The results obtained from the ablation tests are summarized in Appendix Figure A.6. As shown by the tests, the most significant degradation occurs in the distribution and organization of the landscape, as is expected for histogramming methods in latent spaces at high noise (Evans *et al*., 2025). For the decoded conformational states, a similar motion was identified across different noise levels as exemplified by the maps (in blue and red) presented in the figure. However, the amplitude of the motion observed in the maps was slightly lower at high noise levels. This result aligns with the degradation expected from more standard losses such as MSE. Still, the estimated landscape retains a partially straight-line shape even after degradation due to noise and the limited amount of data simulated in the dataset. Given the limited number of images in this simple test (500), it is expected that results will improve as more images become available. From this experiment, we can confirm that noise (at the levels we tested) did not prevent the application of the SW to infer heterogeneity present in a dataset of synthetic images.

## 4 Discussion

The practicality of gradient based methods depends on a confluence of three elements: suitably broad basin of attraction, model specification (avoid bias from miscalibration), and computational feasibility. As shown in Figure 3, if we accurately calibrate the background/contrast, there is a large basin of attraction for gradient-based methods using an OT loss. While it is not computationally feasible to perform many gradient evaluations on the exact Wasserstein loss (e.g. with POT’s `ot. emd2`), instead we can use a inexpensive approximation of the gradient with SW. Notably, our use of the Sliced Cramér–von Mises proxy demonstrates that exact adherence to the Wasserstein metric is not strictly necessary for inference. The proxy preserves the critical spatial signal required to guide gradient descent from a far away initialization, while avoiding the computational cost of quantile inversion. While the SW calculation yields no transport plan, we are not interested in explicitly transporting pixels in the image because we aim to optimize forward model parameters upstream. By leveraging automatic differentiation in cryoJAX, a differentiable forward model simulator, we compute gradients that act on any differentiable parameter, transporting its value iteratively in the direction that minimizes the SW loss. Our work demonstrates that the SW loss provides a broader and smoother optimization landscape than MSE. Both the 3D alignment and 2D single-particle examples show that when the signal is high or the background levels are known, the SW loss landscape is markedly more robust for challenging multi-parameter inference tasks.

This finding suggests a complementary role for the two loss functions: SW provides the wide basin necessary to jointly infer structure and background parameters from distant initializations, while MSE offers an unbiased, albeit narrow, refinement target once the model is within the basin of attraction. This was already noted in the case of 3D rigid body alignment for global alignment of similar but not identical volumes under the 1-Wasserstein distance (Singer & Yang, 2024), and alignment of partially overlapping volumes under the unbalanced Gromov-Wasserstein divergence (Tajmir Riahi *et al*., 2025). In our case, a benefit to the MSE in the basin of attraction is the robustness of high noise levels. In any case, even with an SW loss, these biases could be overcome by averaging over many noisy particles, in the sense that the mean value of the estimated CV over many noisy images is near to the mean ground truth value. However, when inferring other information about the overall distribution of heterogeneity, care would need to be taken in the statistical framing of the problem as emphasized in (Evans *et al*., 2025; Espinosa *et al*., 2025), where high variance of local estimates can overly broaden source distribution estimates from finite measurements.

Background noise calibration is essential for the SW loss. The spiral toy example suggests that a SW failure mode would be in a high noise regime where the local per image SNR *a priori* is in a large interval. In this case, if the underlying estimated projection is sufficiently inaccurate (e.g. incorrect pose, heterogeneity, CTF) then the SW may be driven to very large but unrealistic background, biasing other local variables. This is because with increasing background the normalized images both approach a uniform distribution and thus the SW between them approaches zero. To avoid this, the SW loss could be combined with MSE at different stages, and also the background level could be bounded using prior information—for example by bounding some reasonable value above and below by 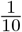 to 10 fold. Local background estimation could be amortized over the dataset, or more formally linked in a hierarchical probabilistic model, where each local background level is drawn from a shared global distribution. Empirical and theoretical work measuring and modeling ice thickness (Rice *et al*., 2018; Peet *et al*., 2019; Armstrong *et al*., 2019) and solvation (Shang & Sigworth, 2012; Parkhurst *et al*., 2024; Himes & Grigorieff, 2021) is a rich source of information for strong background contrast priors. Moreover, recent work analyzed the noise sensitivity of Wasserstein distances under minimal assumptions on multivariate image noise (Lager *et al*., 2025), showing that signed 2-Wasserstein distances enjoy more favorable error scaling than Euclidean norms. While these theoretical results are valuable, our findings highlight the need for further theory addressing (i) sensitivity to calibration bias (unknown per-particle scaling and background) and (ii) how noise shapes the basin of attraction in the OT landscape.

We note that although low-dimensional problems like the spiral example could, in principle, be solved by grid search, recent work has shown that sliced Wasserstein techniques enable efficient rotational searches (Shi *et al*., 2025). The key advantage of using SW-based gradients is that they enable joint, end-to-end optimization of many model parameters simultaneously. In cryo-EM this includes not only local per-image parameters (e.g., pose, CTF, translations, deformations) but also global parameters shared across images (e.g., information about the biomolecular ensemble distribution). In future experiments, we plan to assess the performance of the SW on experimental data by applying our contrast calibration procedure (see Extended Methods section 4.4). We will focus not only on the effect of noise but also on the background inconsistencies introduced by cryo-EM image normalization. This raises important open questions about identifiability and the statistical framing of such optimization (Klindt *et al*., 2024), suggesting future directions involving distributionally robust OT formulations and OT-based manifold optimization.

In conclusion, we propose transport-based losses like the SW and its efficient variants as a computationally feasible workhorse that can be used in place or together with MSE to broaden the basin of attraction for gradient based methods, at acceptable computational cost. Future theoretical work should focus on how to calibrate the contrast, and point out inherent biases in inherent high noise regimes. We hope OT losses like the SW can be applied to challenging flexible specimens that have minimal direct overlap of pixel intensity, to offer statistical knowledge of bimolecular structure and dynamics.

## Author Contributions

GW and ML conceived the project. GW and ML developed code for the goto and goto-swap repositories. DH developed code for results related to Zernike3D. GW, DH and ML performed the analysis and drafted the manuscript. PC and KDD provided supervision. All authors edited the manuscript.

## Acknowledgements

The authors thank Michael J. O’Brien and David Silva-Sánchez for cryoJAX support. The Flatiron Institute is a division of the Simons Foundation.

## Code Availability

Code for reproducing non-cryoJAX results in this manuscript can be found at https://github.com/minhuanli/GOTO. Modular implementations of the SW loss APIs for methods developers can be found at https://github.com/flatironinstitute/GOTO-SWAP, along with experiments using cryoJAX.

## A Extended Methods

### A.1 Forward model and experimental setup

#### A.1.1 CryoJAX simulated data

For the first set of experiments on balancing the tradeoff of accuracy and compute, and performing joint inference of local image parameters, we used cryoJAX to simulate projections of a 9 residue chignolin peptide (PDB id 1UAO), using only non-hydrogen atoms. The image was 128×128 pixels with 0.5 Å per pixel. Noise-free projections were simulated using `cryojax.simulator .make_image_model` with `quantity_mode=‘intensity’`, using a `GaussianMixtureVolume` volume with `amplitudes=1.0, variances=0.25`. For the CTF, we used `ContrastTransferTheory` with `ctf=AstigmaticCTF (defocus_in_angstroms=2000), envelope=FourierGaussian(amplitude=1, b_factor=10)`. Poses were encoded and optimized using `QuaternionPose`. Poisson noise was simulated using the cryoJAX intensity projection, `lambda` as `jax.random.poisson(…, dose*(lambda + background))`, at various dose and background values.

#### A.1.2 CryoJAX forward model for joint inference

For joint inference experiment specifically, we performed gradient descent through the MSE and SW between a reference image (explained in section A.1.1), with defocus 2000 Å, at with coordinates unmodified by pose and a collective variable (CV) which translates coordinates away from the *x* = *y* plane (i.e. translates the *x, y* coordinates tearing the specimen in two pieces). We optimized defocus, *t*_*x*_ and *t*_*y*_ translation, rotation, and CV. The optimization variables are initialized at 3000Å defocus, *t*_*x*_ = +5 Å, *t*_*y*_ = +5 Å, and ZXZ Euler angles (*ϕ* = 5, *θ* = 5, *ψ* = 5) degrees, and *CV* = 5. We used `cryojax.simulator.QuaternionPose(wxyz=…)` to encode the rotation during inference. Every gradient step the optimization variables were used to simulate through the cryoJAX forward model, and updated by a gradient with respect to the MSE or SW loss. In order to account for physical units, we used the diagonal preconditioners in Table 2. We determined these values by performing inference on one parameter at a time, holding the others fixed, and adjusting the step size so such that convergence was reached with the same amount of steps.

#### A.1.3 PyTorch forward model

For experiments that did not involve the CTF, but only projected 3D coordinates onto the plane (i.e. 3D rotational alignment, and spiral deformation), we used a PyTorch forward model. This differentiable forward model projected coordinates directly to 2D image using a sparse multi-index data structure in PyTorch `torch.sparse_coo_tensor`. In brief, coordinates were parametrically deformed (e.g. rotated and deformed by the collective variable) and a blob of intensity was generated at their centre with a 2D Gaussian bump function. We assumed a fixed amplitude and standard deviation (std), and truncated at 3 std away.

#### A.1.4 Helical spiral experiment setup

We parametrized the helical spiral using an angular parameter

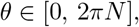

where *N* = 10 is the prescribed number of full revolutions. Given a total number of sample points *M* = 2000, we discretized this interval uniformly:

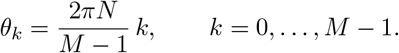

Let *r >* 0 denote the (constant) radius of the spiral. We embeded the curve in ℝ^3^by allowing its axial coordinate to increase linearly with *θ*, while the transverse coordinates follow a circular trajectory. Specifically, we define

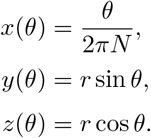

The resulting parametric curve is therefore

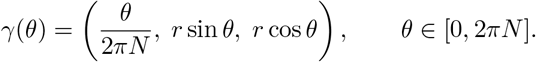

After sampling the points

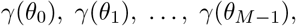

we recenter the cloud by subtracting its empirical mean,

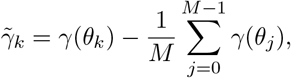

so that the final point set

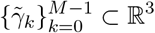

is centered at the origin.

#### A.1.5 Contrast Calibration

For the noise experiment, we generated a noisy image of our projected spiral *s*(*CV* =4) = *s*_gt_ (whose intensity values ranged from 0–2.5 on a *N* = 64^2^ pixel image) at a dose *d*_gt_ = 30 and background *b*_gt_ = 100, and sampled with Poisson noise:

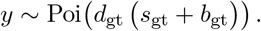

The resulting image was normalized by Z-scoring,

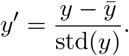

For each candidate pair of collective variable (CV) and background, we generated a clean forward model image

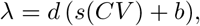

and then back-transformed the normalized noisy image *y*′ into the corresponding standardized Poisson domain of *λ* = (*λ*_1_, …, *λ*_*N*_), where each *λ*_*i*_ ∈ ℝ_≥0_ is the pixel-wise value of the noiseless calibrated simulation. For a Poisson random vector with parameter *λ* = (*λ*_1_, …, *λ*_*N*_), the population mean and variance of the corresponding mean and variance for Z-scoring satisfy

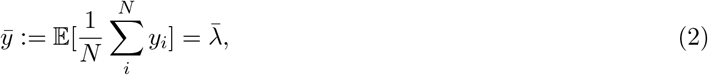

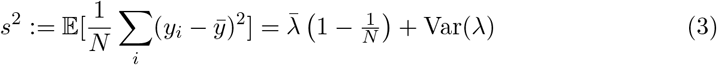

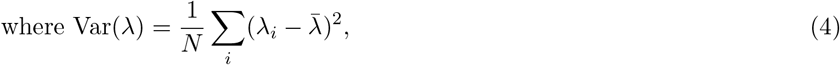

which gives the expected sample mean 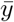 and sample (population-form) variance *s*^2^ for the Poisson-distributed image.

Thus the back-transformation mapping the normalized noisy image *y*′ to the Poisson domain of a candidate image is

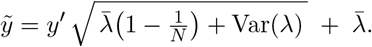

This ensures that the normalized noisy observation is compared fairly against each CV/background hypothesis under the correct Poisson statistics induced by *d*(*s* + *b*).

### A.2 KL Divergence Between Poisson Distributions

Let

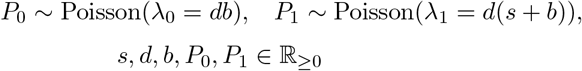

We are interested in computing the KL divergence,

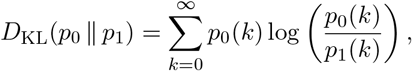

where *p*_*i*_ is the PMF of the Poisson distribution:

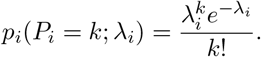

Substituting into KL:

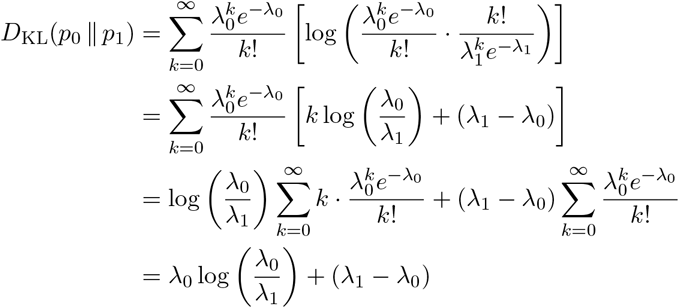

And finally substituting *λ*_0_ = *db, λ*_1_ = *d*(*s* + *b*):

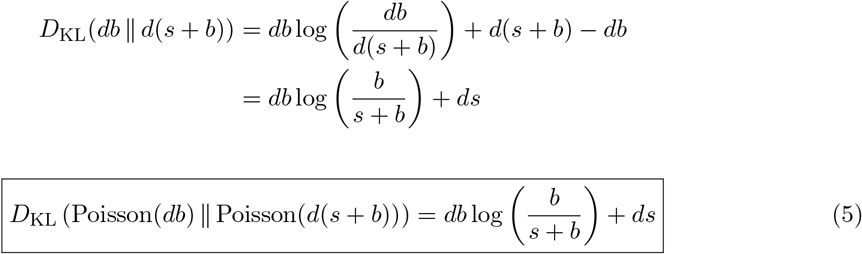

The correspondence to the signal to noise ratio of Gaussian white noise images is shown in Figure A.1, which can give a “ball-park” correspondence between our experiments and the many others that use a Gaussian white noise model. For an image’s *D*_KL_, the value of *s* and hence *λ*_*i*_ is different in each pixel, and so we averaged the *D*_KL_ across pixels instead of averaging *s* before computing the *D*_KL_. Note that matching *σ*^2^ = Var[Poi(*aλ* + *b*)] = *d* · 𝔼[*λ*] + *b*, and plotting “Gaussian SNR” vs. “Poisson SNR” gives a linear relationship with a slope not equal to one, but similar visual appearance.

### A.3 2D Earth Mover’s Distance Implementation Details

For the exact OT baseline, the input images of size *n* × *n* were flattened into vectors of length *n*^2^ and normalized to sum to unity, ensuring they represent valid probability measures. The cost matrix *C* was constructed as the squared Euclidean distance between pixel coordinates, resulting in a matrix of size *n*^2^ × *n*^2^. To ensure numerical stability across different backends, we normalized the cost matrix by dividing by its maximum element: *C* ← *C/* max(*C*). This scaling is linear and therefore does not alter the optimal transport plan.

We utilized the Python Optimal Transport (POT) library’s `ot.emd2` function, which employs a network simplex algorithm. To handle complex transport problems where the default maximum solver iterations might be insufficient, we used 10^6^ from the default 10^5^. Unlike entropy-regularized approaches (e.g., Sinkhorn), this exact solver does not require a regularization hyperparameter, though convergence is subject to standard floating-point precision tolerances.

Crucially, the computational complexity of the network simplex algorithm scales cubically with the number of pixels *O*((*n*^2^)^3^), rendering it impractical for direct unrolling in a computation graph. We therefore utilized the analytic gradient hooks provided by POT for JAX and PyTorch. These hooks treat the solver as a black box during the forward pass; during the backward pass, they compute gradients with respect to the marginals and cost matrix using the analytical solution derived from the optimal transport plan (via the envelope theorem), rather than differentiating through the solver’s internal operations. The envelope theorem (Milgrom & Segal, 2002) states that the gradient of an optimization problem’s optimal value with respect to its parameters depends only on the objective evaluated at the optimal solution, allowing gradients to be computed without differentiating through the solver itself.

### A.4 Sliced Wasserstein Implementation Details

#### Sparse Projections (Greedy Matching, *SW*_GM_)

For the coordinate-based method, we ported the sorted 1D EMD algorithm (specifically `ot.lp.emd_wrap.emd_1d_sorted`) from the POT library’s Cython implementation to native JAX and PyTorch. The algorithm sorts the marginals by their 1D spatial location and performs an iterative greedy mass transfer without backtracking.

- **Differentiable Sorting:** We sorted the spatial coordinates using standard differentiable sort operations. The resulting indices were used to reorder the space and marginal, ensuring gradients propagate through the coordinates and weights while the gradient do not need to pass through the values of the sorting indices.
- **JAX Implementation:** To enable JIT compilation, we replaced the standard while loop with `jax.lax.scan`. We set the maximum loop iterations to the sum of the lengths of the two input vectors, which is the theoretical upper bound for the greedy match.
- **PyTorch Implementation:** We utilized a standard Python while loop, which is natively supported by PyTorch’s autograd engine.
- **Parameters:** We used a numerical tolerance of 10^−12^ to determine mass equality during the iterative assignment.

#### Dense Projections (*SW*_iCDF_, *SW*_CDF_, and *SW*_GM_)

For the Radon transform-based method, image rotation was performed using `dm_pix.rotate(…, mode=‘wrap’)` (JAX) or `torch.nn.functional .grid_sample(…, mode=‘nearest’, padding-mode=‘circular’, align-corners=False)` (PyTorch). To compute the 1D distance:

- **iCDF Integration:** We interpolated the inverted CDF onto a regular grid of 1000 points using `searchsorted` with default options for JAX and PyTorch and computed the integral via trapezoidal quadrature.
- **CDF Computation:** We computed the CDF via a normalized cumulative sum. To ensure numerical stability when normalizing (avoiding division by zero), we added a small number to the denominator at machine-precision (`jnp.finfo(input_dtype).eps, torch.finfo (input_dtype).eps`), dependent on the data type of the input.
- **GM:** as above

### A.5 3D Alignment Benchmark Details

We evaluated the 3D alignment performance on a set of 13 biomolecular structures obtained from the Electron Microscopy Data Bank (EMDB) and previously used for 3D alignment (Riahi *et al*., 2023). The specific entries used were: EMD-1717, EMD-2677, EMD-3240, EMD-3341, EMD-3342, EMD-4547, EMD-6284, EMD-6475, EMD-8724, EMD-8764, EMD-8881, EMD-9515, and EMD-38142.

For each system, we generated a benchmark set of 100 initial poses by applying independent random rotations to the ground truth volume. These rotations were drawn uniformly up to a cut-off of 90 degrees geodesic distance. We then ran the optimization for 2000 gradient steps using either *SW*_GM_ or MSE loss and recorded the final rotation error (geodesic distance to ground truth). For each *SW*_GM_ loss calculation, we set 12 random projections, with 12 random pixels in each projection. Note that the the two volumes shared the same grid and random pixels.

To quantify robustness for a single system, we computed the cumulative distribution function (CDF) of these 100 final rotation errors. We then calculated the Area Under the Curve (AUC) of this CDF, normalized such that an AUC of 1.0 corresponds to all 100 runs converging to 0 deg error, while lower values indicate that a fraction of the runs remained trapped in local minima with large residual errors. The distributions shown in Figure 2b consist of these per-system AUC values.

### A.6 Zernike3D

Zernike3D (Herreros *et al*., 2023) is an approach previously introduced in the field to estimate diverse structural states from a set of particle images in a CryoEM dataset. Starting from a 3D reference state (e.g., a consensus volume), the method estimates the vector field warped the reference geometry towards a target state captured in a 2D image. Instead of directly optimizing the complete vector field, the approach is to expand the field in terms of the Zernike3D basis as follows:

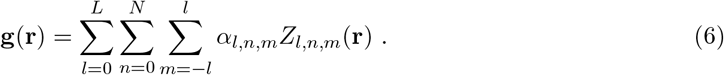

Where **g** is the vector field, **r** is a given position in the reference state, *α*_*l,n,m*_ is a set of Zernike coefficients, and *Z*_*l,n,m*_ is a given component of the Zernike3D basis.

From the previous equation, it is evident that estimating the vector field reduces to finding the set of coefficients *α*_*l,n,m*_ used to expand the desired vector field. We can express the previous optimization as:

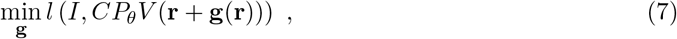

where *I* represents an experimental image in a dataset, *V* is the reference state, *C* the CTF estimated for that particle, *P*_*θ*_ the projection operator along the 3D direction and in-plane shift defined by the parameters *θ*, and *l* is a loss function comparing the experimental image with the theoretical projection of the warped reference state.

For the experiments presented in this work, we compare the well-known Mean Squared Error (MSE) loss function against the Sliced Wasserstein Distance (SWD). In both cases, the coefficient optimization follows the scheme outlined in the previous equation, with the loss function as the only notable difference.

**Figure A.1:**
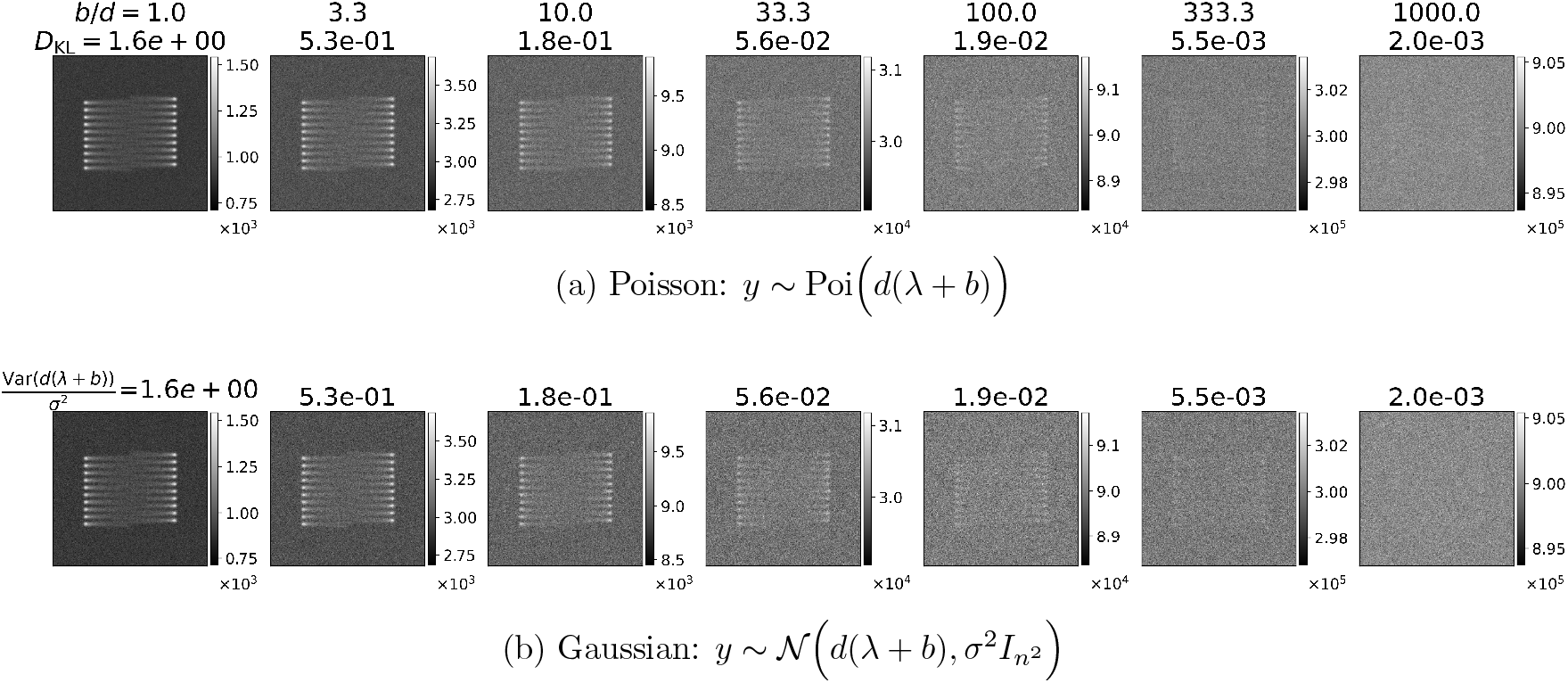
Poisson and Gaussian SNR. For comparison, we show example **(a)** Poisson images (dose *d* = 30 and background *b* ∈ {30, 10^2^, 3 · 10^2^, 10^3^, 3 · 10^3^, 10^4^, 3 · 10^4^}) next to the **(b)** Gaussian image at the “same” SNR. For “Poisson SNR”, we use the KL definition in Eq. (5), where we average *D*_KL_ across each value of the signal, which is different from averaging the signal first and then computing *D*_KL_. For “Gaussian SNR”, we use the variance *σ*^2^ that gives the same SNR, according to the typical SNR definition of the ratio of signal variance to noise variance: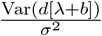, where *λ*_*i*_ is the value of *i*th pixel before scaling by the dose and background.

## B Sliced Wasserstein Accuracy

### B.1 Accuracy: Data Generation Process

We used the data generation process outlined in Algorithms 1, 3, and 2 as numerical tests to verify the numerical equivalency between SW and EMD, in terms of averages, scaling by the dimension, and square/square root definitions (e.g. *W*_1_ versus *W*_2_).

#### Algorithm 1

Monte Carlo comparison between 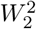 and Sliced Wasserstein

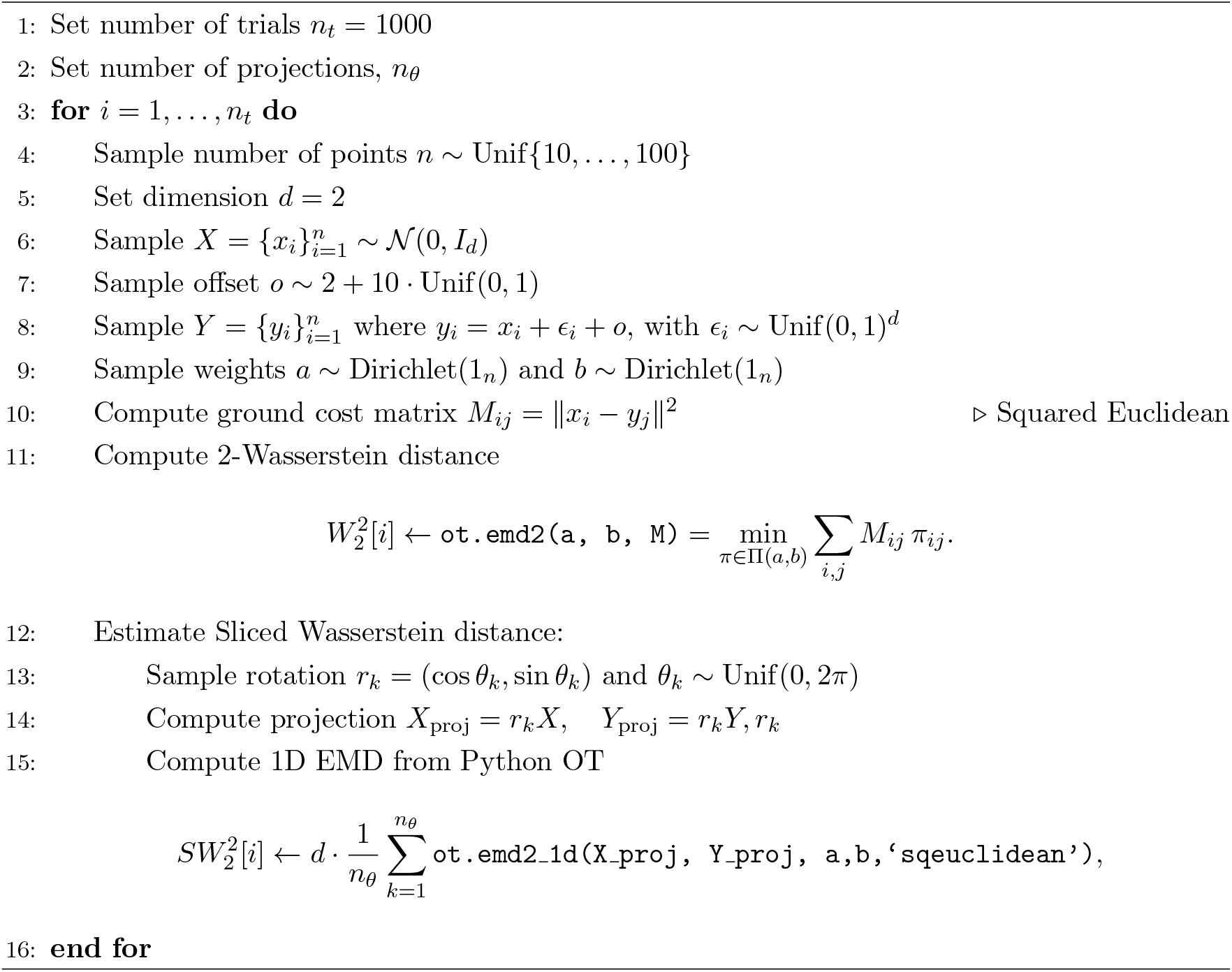

#### Algorithm 2

Quantitative comparison between inverse-CDF Wasserstein estimator and 1D EMD

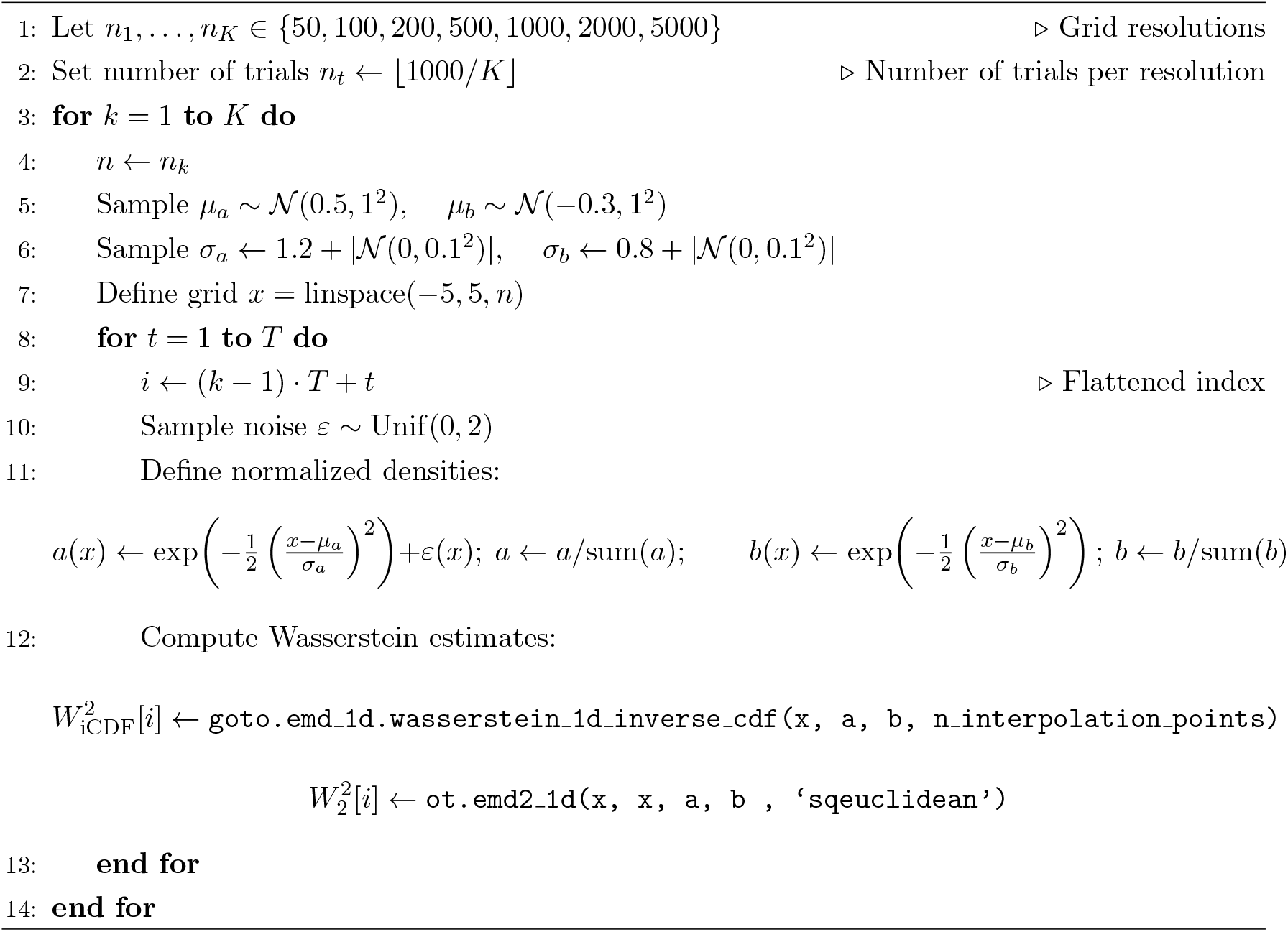

#### Algorithm 3

Monte Carlo comparison of 1D OT methods

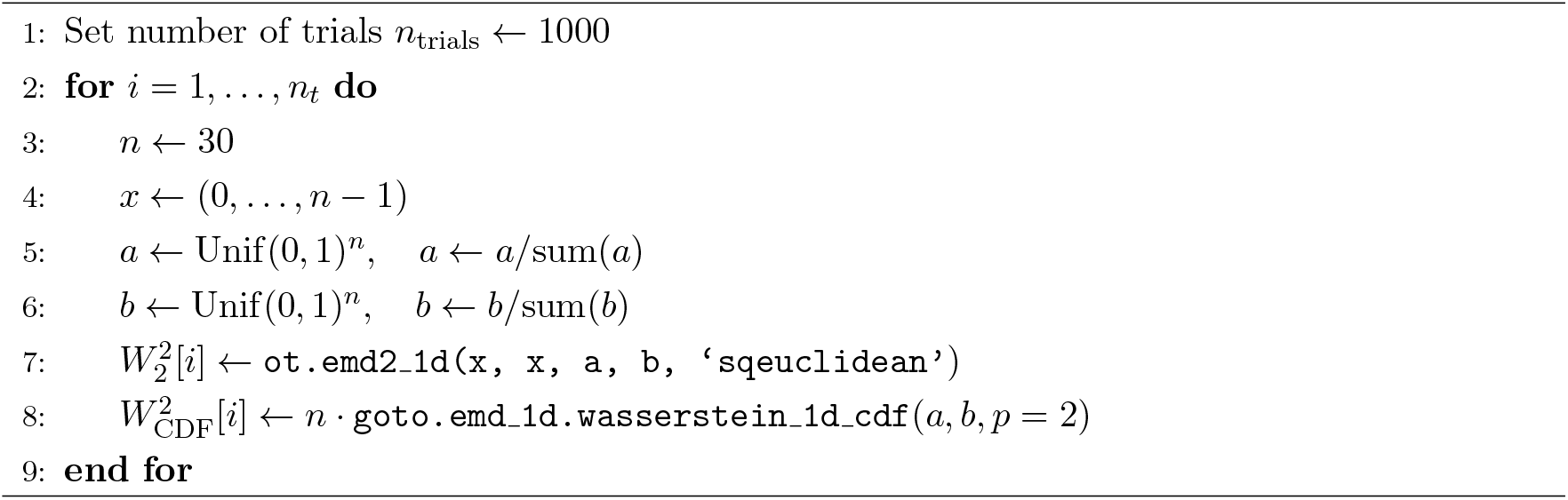

In brief, we compared *W*_2_ to *SW*_2_ (via averaging over 1D projections) through the POT implementations of `ot.emd` and `ot.emd_1d`, which wraps the Greedy Matching algorithm in `ot.lp.emd_wrap .emd_1d_sorted`. Then we verified our differentiable reimplementation of `ot.lp.emd_1d`. We then matched this 1D EMD to the iCDF and CDF, thereby linking 2D *W*_2_ to our three implementations of SW. Figure A.2 shows plots of the pairs 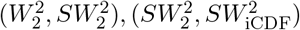, and 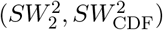.

Figure A.3 shows accuracy as a function of the number of projections.

## C Noise bias of SW

We repeated the experiment in section 3.3 for different random draws of Poisson shot noise. Figure A.4 shows that SW under and over estimates the ground truth CV and background, while MSE is more robust to noise. Table 3 quantifies the bias. We also used different values of dose to change the SNR. As the SNR increases (increasing dose), the variance of the inferred global minima decreases. The min/max values show that the inferred global minima does not reach the boundary for the region on which we evaluated the landscape.

**Table 3:** Landscape global minima statistics across noise trials. Summary statistics are over 100 trials, at a background (b) grid spacing of 50-200 in increments of 3, and a CV grid spacing of 1-8 in increments of 0.14. The ground truth values are CV=4, and b=100. The SW was computed via SW_CDF_ with 100 projections.

| dose | Loss | SNR (nats) | CV (mean ± std) | CV (min, max) | b (mean ± std) | b (min, max) |
| --- | --- | --- | --- | --- | --- | --- |
| 3 | SW | $6.30 \times 10^{-3}$ | $4.27 \pm 0.76$ | (3.00, 6.57) | $98.43 \pm 24.94$ | (53.06, 166.33) |
| | MSE | | $4.00 \pm 0.00$ | (4.00, 4.00) | $101.34 \pm 7.56$ | (83.67, 123.47) |
| 10 | SW | $2.10 \times 10^{-2}$ | $4.07 \pm 0.36$ | (3.14, 5.00) | $99.32 \pm 13.90$ | (65.31, 141.84) |
| | MSE | | $4.00 \pm 0.00$ | (4.00, 4.00) | $100.27 \pm 4.27$ | (92.86, 111.22) |
| 30 | SW | $6.30 \times 10^{-2}$ | $4.00 \pm 0.19$ | (3.57, 4.57) | $101.28 \pm 7.57$ | (86.73, 123.47) |
| | MSE | | $4.00 \pm 0.00$ | (4.00, 4.00) | $99.81 \pm 2.34$ | (89.80, 105.10) |
| 100 | SW | $2.10 \times 10^{-1}$ | $4.01 \pm 0.10$ | (3.71, 4.29) | $100.11 \pm 4.26$ | (92.86, 111.22) |
| | MSE | | $4.00 \pm 0.00$ | (4.00, 4.00) | $100.14 \pm 1.49$ | (98.98, 102.04) |

## D Zernike3D rank based heterogeneity accuracy

Although the previous comparisons show that SWD yields a more accurate representation of the expected landscape, it is possible to find other approaches to quantitatively assess how close the estimated Zernike3D are to the ground-truth NMA vectors. Therefore, we propose using the information imbalance metric, which is expressed as:

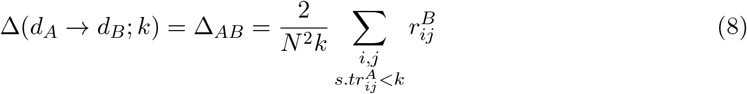

The results obtained from the information imbalance analysis are shown in A.5. For each metric (i.e., SWD and MSE), five independent executions of Zernike3D were performed to assess execution stability. As shown in the plot, the SWD landscapes yield an information imbalance value closer to the origin, indicating they are closer to the NMA ground truth. Our findings suggest that SWD is a promising loss function candidate with the potential to achieve better accuracy than other well-established metrics, such as MSE.

**Figure A.2:**
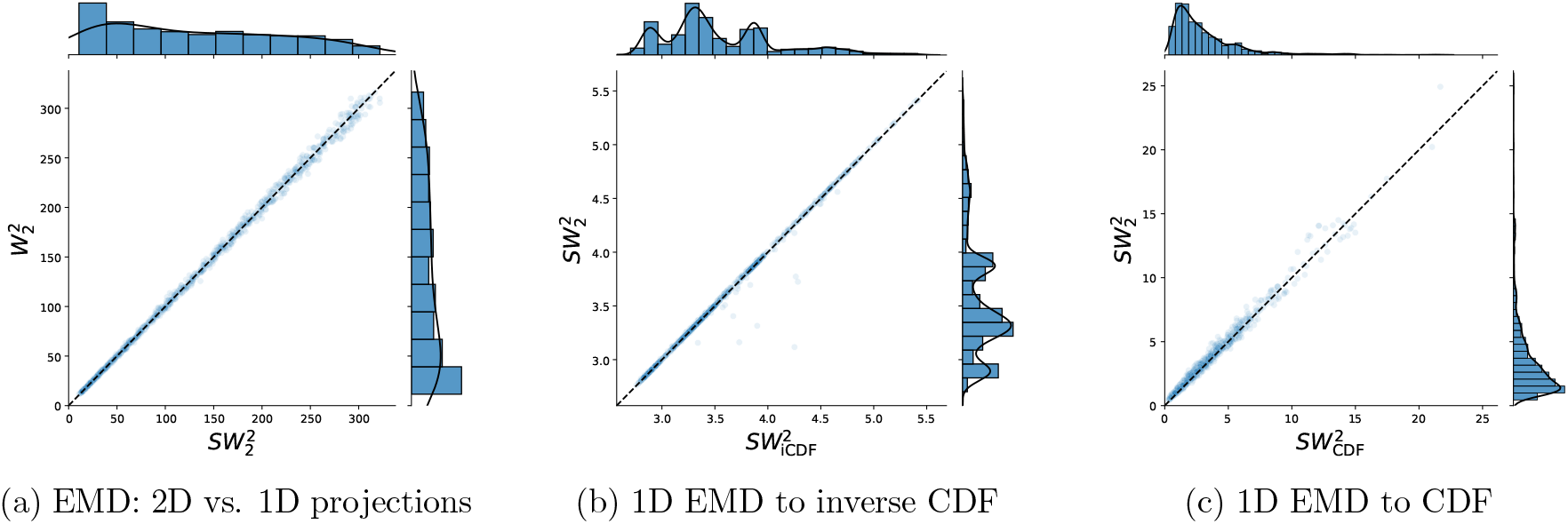
Accuracy. **(a)** We benchmarked the accuracy of 2D 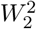 to 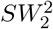 (the average 1D 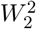) over a set of *n*_*θ*_ = 1000 uniformly random projections, both implemented in Python OT. The data generation process is Algorithm 1. **(b)** We benchmarked 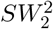 against the 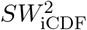. The data generation process if Algorithm 2. **(c)** We benchmarked 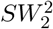 against 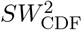. The data generation process is Algorithm 3.

**Figure A.3:**
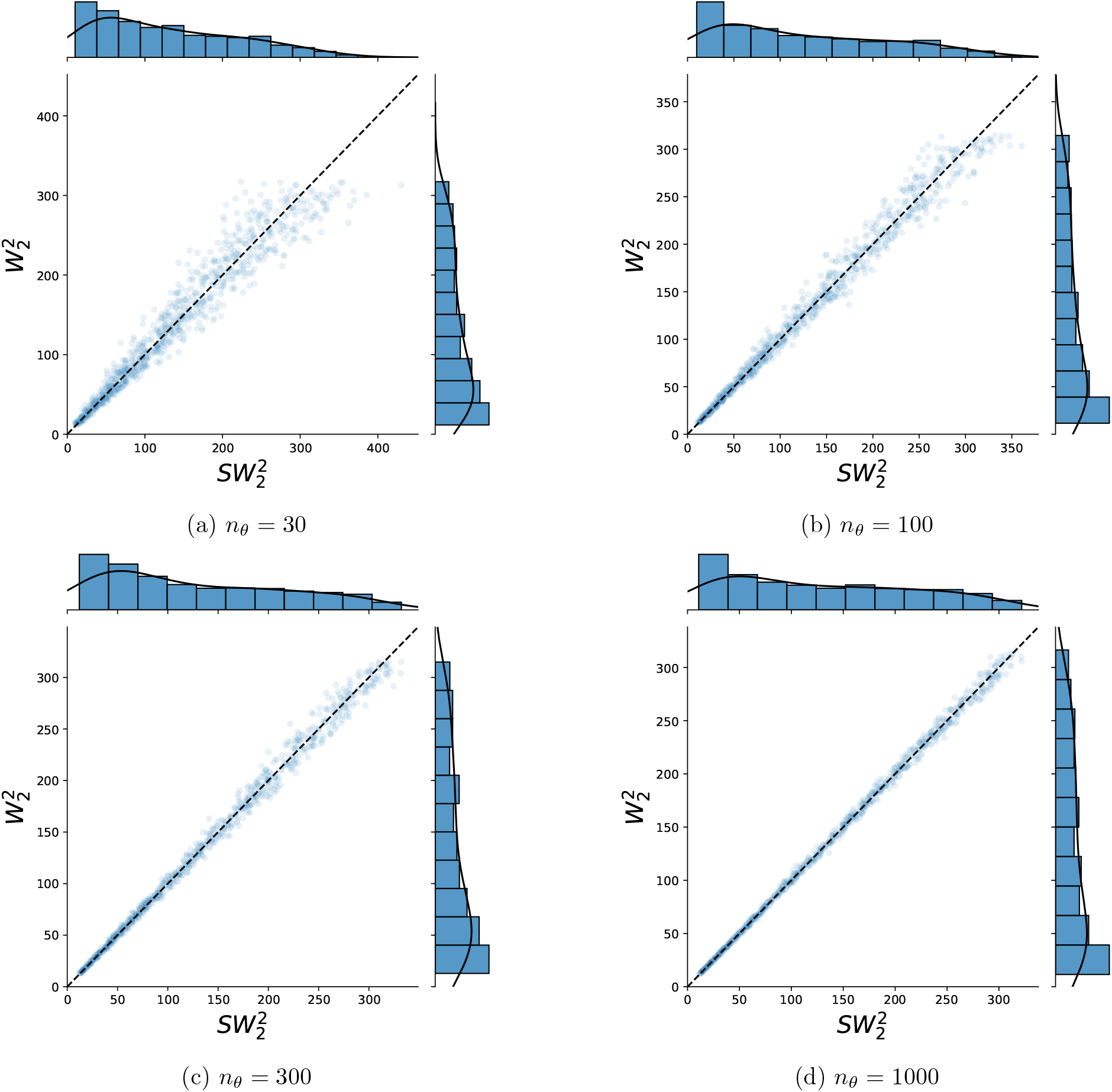
Accuracy. We benchmarked the accuracy of 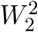 to 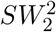, the average 1D 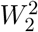, over several sets of *n*_*θ*_ = {30, 100, 300, 1000} uniformly random projections, both implemented in Python OT. The data generation process is Algorithm 1.

**Figure A.4:**
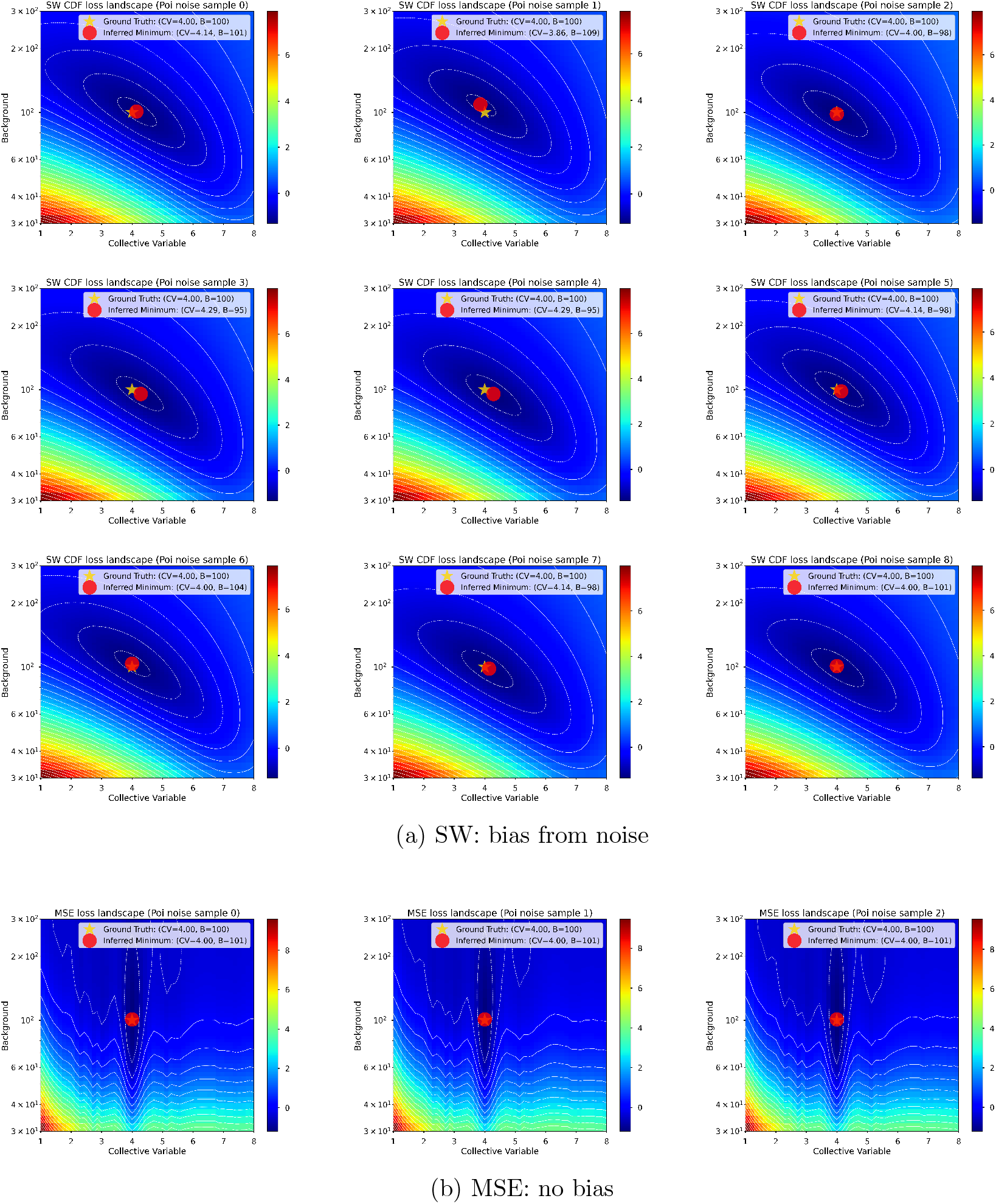
Sliced Wasserstein optimal parameters are biased by noise. **(a)** Each SW landscape corresponds to a different noise instantiation. The inferred collective, CV, and background level, *b*, are around the ground truth value – sometimes higher, sometimes lower. **(b)** The MSE global minima contains to bias from the ground truth *b* and CV values.

**Figure A.5:**
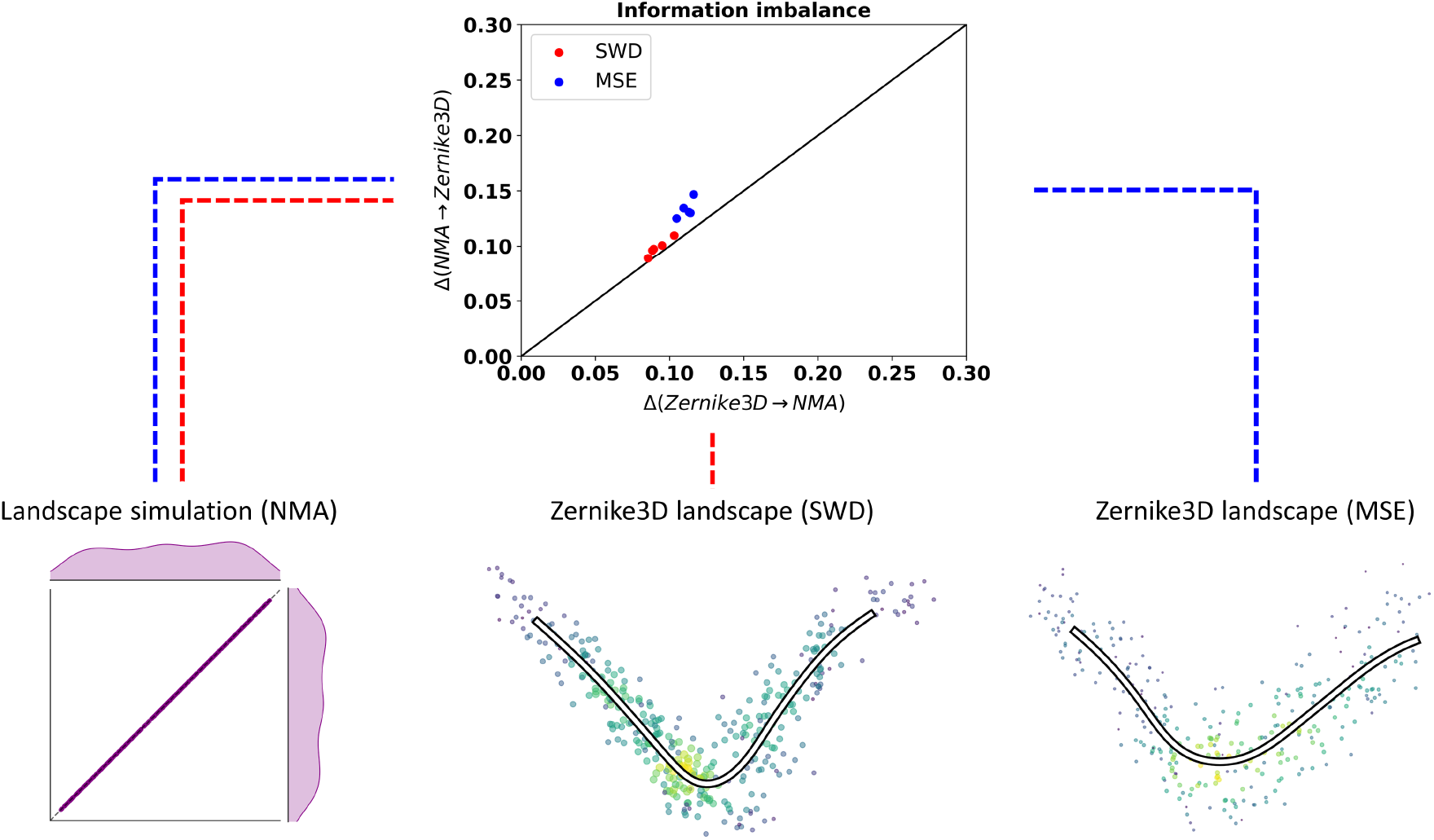
Zernike3D analysis comparing NMA against SWD and MSE with the information imbalance metric. Information imbalance tests comparing the original NMA vectors against the Zernike3D landscapes. The results show that the SWD metric yields a landscape closer to the ground truth with improved stability compared to MSE.

**Figure A.6:**
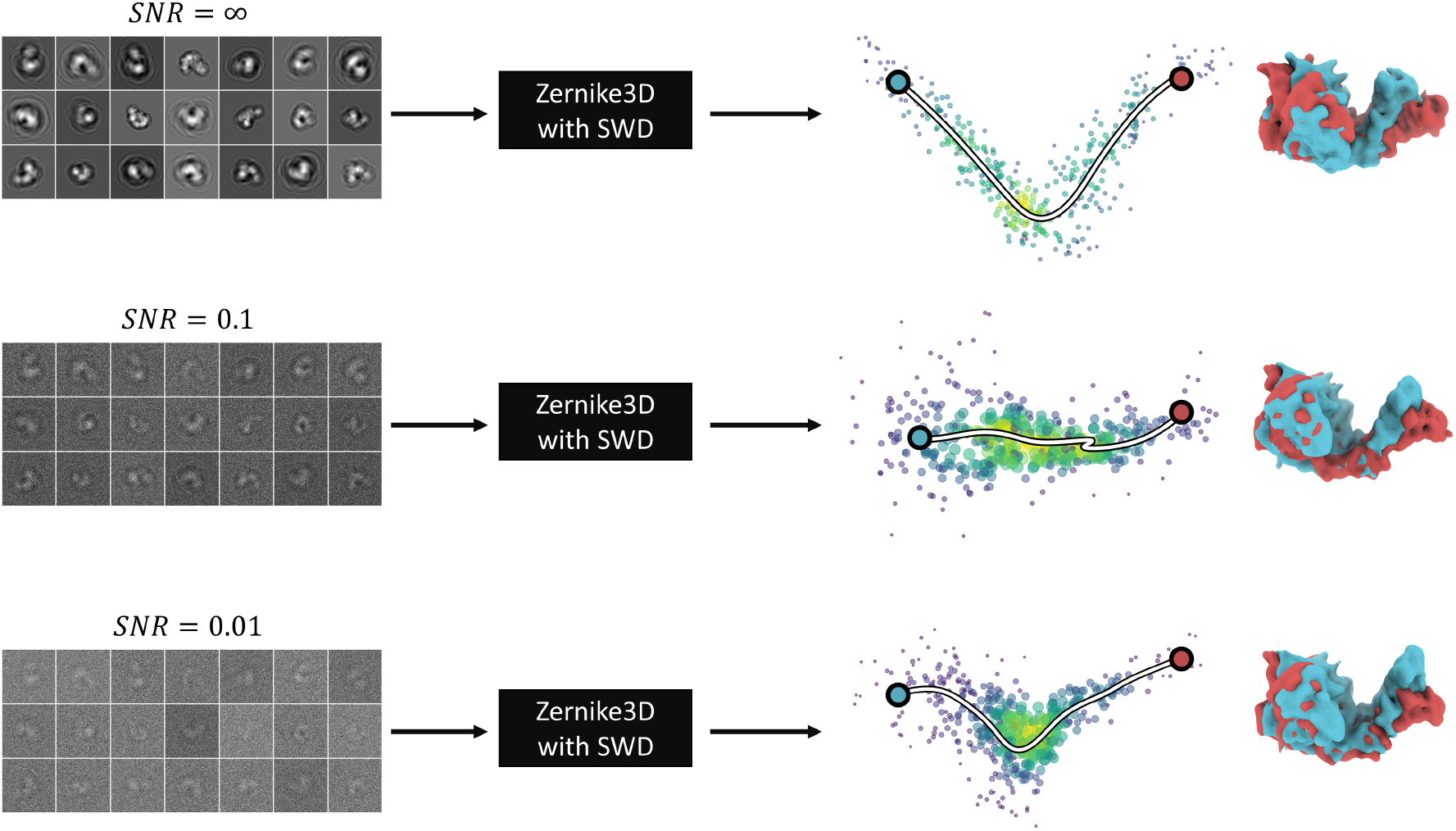
Zernike3D noise ablation test. Results obtained from the noise ablation test with Zernike3D and the SWD. The results show that the most significant degradation occurs in landscape organization, while the detected conformational changes remain more stable.

